# Metataxonomic review to elucidate the role of the microbiome in Celiac disease across the gastrointestinal tract

**DOI:** 10.1101/2021.08.08.455560

**Authors:** Juliana Arcila, Viviana Loria-Kohen, Ana Ramírez de Molina, Enrique Carrillo de Santa Pau, Laura Judith Marcos-Zambrano

## Abstract

**Background:** Dysbiosis of the microbiome has been related to the Celiac disease (CeD) progress, an autoimmune disease characterised by gluten intolerance developed in genetically susceptible individuals under certain environmental factors. The microbiome contributes to CeD pathophysiology modulating the immune response by the action of short-chain fatty acids (SCFA), affecting gut barrier integrity allowing the entrance of gluten derived proteins, and degrading immunogenic peptides of gluten through endoprolyl peptidase enzymes. We reviewed state of the art in taxonomic composition for CeD and compiled the larger dataset of 16S prokaryotic ribosomal RNA (rRNA) gene high-throughput sequencing for consensus profiling. We present for the first time an integrative analysis of metataxonomic data from CeD patients, including samples from different body sites (saliva, pharynx, duodenum, and stool). We found the presence of coordinated changes through the gastrointestinal tract characterised by an increase in *Actinobacteria* species in the upper tract (pharynx and duodenum), and an increase in *Proteobacteria* in the lower tract (duodenum and stool), as well as site-specific changes evidencing a dysbiosis in CeD patients’ microbiota. Moreover, we described the effect of adherence to a gluten-free diet (GFD) evidenced by an increase in beneficial bacteria and a decrease in some *Betaproteobacteriales* but not fully restoring CeD-related dysbiosis. We illustrate that the gut microbiota acts as an enhancer of immune response in CeD through the production of lipopolysaccharides and other bacterial components that activate the immune response and by decrease SCFA producers bacteria. Furthermore, microbial changes observed through the gastrointestinal tract of CeD patients may help manage the disease and follow-up GFD treatment.

## 1. Background

The gut microbiome comprises the genome of all the microorganisms that inhabit the gut, and it plays a crucial role in nutritional, metabolic, physiological and immunological processes of the body; it also produces several metabolites such as short-chain fatty acids (SCFAs), branched-chain amino acids, vitamins, and hormones, that affects several metabolic and inflammatory pathways [1] Numerous immune-mediated diseases have been linked to shifts in the gut microbiome composition, including celiac disease (CeD) [2].

CeD is an autoimmune disease affecting the small intestine, ranging from an intraepithelial lymphocytosis to the total atrophy of intestinal villi as a response to gluten consumption [3]. This disease is characterised by a genetic predisposition given by the alleles coding for the HLA-DQ2 and/or HLA-DQ8, the presence of antibodies against transglutaminase type 2 (TG2) and IgA and IgG anti-gluten as well as gastrointestinal (GI) symptoms when consuming gluten-containing foods [3]. The HLA variant with a higher association with CeD is HLA-DQ2, being present in more than 90% of patients. About half of the remaining patients possess an HLA allele coding for HLA-DQ8. However, a small fraction of individuals at genetic risk develops CeD. Suggesting that besides genetic factors, the risk of disease can also depend on non- gluten environmental factors [4].

Studies profiling the microbiome in CeD supports an alteration in the intestinal microbiome of patients suffering from the disease [2]. The intestinal microbiome in CeD comprises an increased number of opportunistic bacteria clades while beneficial clades are decreased [5]. This phenomenon is called dysbiosis and has become subject to active and growing research; as one of the non-gluten environmental factors contributing to CeD development [6–8].

Gliadin is the gluten-derived protein that triggers proinflammatory cytokines in CeD patients, among them IL-6, which stimulates the growth and differentiation of T cytotoxic cells that mediate the destruction of surface epithelial cells [9–11]. Gliadin peptides present multiple proline and glutamine amino acids residues, making them resistant to proteolysis through gastric, pancreatic, and intestinal proteases. This results in incomplete digestion of gluten and the generation of high molecular weight oligopeptides, some of which have immunogenic and toxic properties relevant to CeD. The most studied of these peptides is 33-mer [12, 13].

Recent research has shown that peptidases from different sources can degrade gluten and gluten-derived peptides [14, 15]. In this regard, several bacteria from the human digestive tract (i.e: *Bifidobacterium* spp., *Lactobacillus* spp., *Rothia* spp) can potentially degrade gluten, and a healthy microbiome composition could modulate the symptoms of gluten-related diseases [15]. The presence of several microorganisms that produce proteases able to hydrolyse peptides rich in proline and glutamine residues has been described in the oral cavity, duodenum, and faeces [14, 15].

On the other hand, the tissular Transglutaminase (tTG) enzyme has an essential role in CeD pathogenesis by enhancing gliadin epitopes immunogenicity. This enzyme effectuates specific and organised deamination of high molecular weight derived gliadin peptides giving them an improved affinity to HLA-DQ molecules associated with CeD and forming kinetically stable HLA-DQ-peptide complexes [16]. The HLA-DQ heterodimers are HLA class II molecules expressed on the surface of antigen presenter molecules (APC). In CeD pathogenesis, HLA-DQ molecules associated with the disease play a role in presenting gliadin peptides to T CD4+ cells restricted to recognise those HLA-DQ heterodimers. These specific T cells are present in CeD patients but not in healthy people [9, 17]. Some studies have found that HLA-DQ genotype affects early gut microbiota composition, favouring the colonisation with pathogenic bacteria [18].

The gut microbiome could play a role in triggering CeD in genetically susceptible persons in several ways. First, dysbiosis could lead to an alteration in the intestinal barrier, sustained inflammation or infection can lead to deregulation in the expression of adhesion molecules at tight junctions leading to the entry of microbes and toxic substances and facilitate entry of incompletely digested gliadin peptides in lamina propria [2, 5, 8]. Of several proteins involved in tight junctions, disassembly of zonulin has been implicated in CeD patients [19] gluten peptides and some enteric bacteria, like *Escherichia coli,* can induce this protein [18].

Gliadin peptides and gut dysbiosis can activate innate and adaptive immune systems similarly [20]. Pathogenic gram-negative bacteria activate the innate immune system by activating Toll-Like Receptors (TLR-4), and CD14 complexes recognise bacterial endotoxins and lipopolysaccharide and activate the innate immune system to release proinflammatory cytokines [2, 20]. In patients with CeD, gluten intake is associated with activating gluten-specific CD4^+^ T cells in the lamina propria and upregulation of IL-15, a proinflammatory cytokine [21]. Moreover, gut microbiota can also activate Th1, Th2 and Th17 mediated immune responses similar to upregulation by gliadin peptides [22].

Finally, as discussed above, dysbiosis may also increase the amount and size of gliadin peptides due to differential peptidolytic activity of the gut microbiota [14, 15]. These studies suggest that the gut microbiota affects gluten digestion, intestinal permeability, and the host immune system, all the mechanisms involved in the pathogenesis of CeD.

A schematic representation of the microbiome’s involvement in CeD suggested by the current knowledge in the topic is shown in Figure 1. In the intestinal lumen are the peptides resulting from the degradation of the diet components, including gluten; there is also the microbiome composed of beneficial and opportunistic commensal microorganisms (step 1, Fig. 1). A dysbiosis characterises CeD microbiome with an increase of opportunistic bacteria (mainly from the phylum *Proteobacteria*), these bacteria compete to adhere to the epithelium, and when they reach, they trigger an inflammatory immune response that increases the permeability of the epithelial barrier, allowing the passage of immunogenic gluten peptides to the lamina propria (step 2, Fig. 1). In there, the gliadin peptides are modified by TG2, improving their junction to HLA DQ2/8 heterodimers (step 3, Fig. 1). These modified peptides are delivered by APCs to T lymphocytes (step 4, Fig. 1), producing a chain of immune responses that ends up in the destruction of enterocytes (step 5, Fig. 1) and the production of anti-gliadin and anti-TG2 antibodies (step 6, Fig. 1)

**Figure 1.**
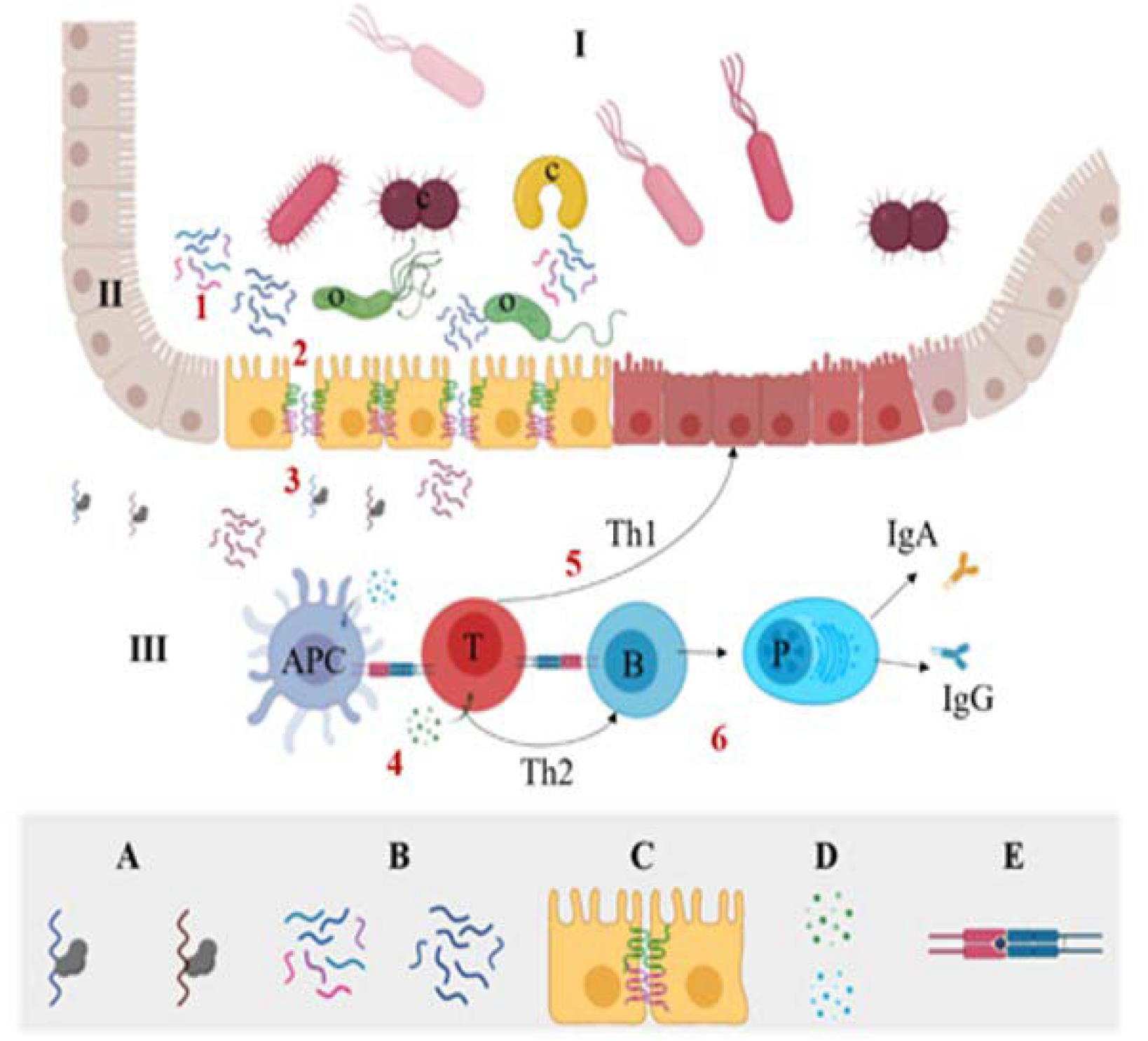
Schematic representation of the sequential process of interaction between gluten, the microbiome and the immune response in the intestine, which leads to the destruction of enterocytes in the pathogenesis of CeD suggested by the evidence to date. I = intestinal lumen; II = epithelium; III = lamina propria; APC = antigen presenting cell; B = cell B; P = plasma cell, o = opportunistic or pathogenic commensal bacteria; c = beneficial commensal bacteria, T = CD4 + T cell. A. tTG-gluten peptide complex: deaminated (right brown), unmodified (left blue), Diet degradation peptides: gliadin peptides (right blue), other dietary peptides (various colors left). C. Tight junctions between epithelial cells: claudin (green), occludin (pink). D. Cytokines: pro-inflammatory (blue), and anti-inflammatory (green). E. HLA DQ2 / 8 heterodimer (blue), immunogenic gliadin peptide (blue circle), TCR (pink).

### 1.1 Gluten-free diet and microbiome

Strict adherence to a Gluten-Free Diet (GFD) and a lifelong exclusion of gluten from the diet is the first-line treatment and is currently the only effective therapy for CeD [23]. A GFD requires the complete exclusion of gluten, a protein complex present in food products from wheat, rye, barley, oats, among other cereals. It comprises only naturally gluten-free food products (e.g., legumes, fruit and vegetables, unprocessed meat, fish, eggs and dairy products) and/or substitutes of wheat-based foods, specially manufactured without gluten or having a gluten content lower than 20 ppm, as per European legislation [24].

GFD has become a trend in contemporary history being associated with an increase in energy and boosting health; however, evidence suggests that GFD is an unbalanced diet with multiple nutritional deficiencies [25]. Gluten restricted products are often associated with an inadequate nutritional value, characterised by a higher fat intake, less vegetal-protein intake, and higher carbohydrate and sugar consumption [24, 25]. Although GFD can reduce the symptoms of CeD in most patients, it does not entirely restore the gut microbiota to that of healthy individuals; moreover, it has been reported that up to 30% of patients will exhibit non-responsive CeD, a condition characterised by the persistent enteropathy and CeD related symptoms after one year on a GFD [26].

### 1.2 Microbiome research and CeD

Traditional methodologies like PCR-DGGE [27, 28] PCR-TTGE [29], cloning [30] and culture [31, 32] for subsequent identification by Sanger sequencing were used to study the microbial composition of the gastrointestinal tract of CeD patients before 16S rRNA gene-based metataxonomic approach become popular. These previous studies set the basis of current knowledge in CeD by revealing a lower microbial diversity in CeD patients with gastrointestinal symptoms than in patients with extraintestinal symptoms suggesting for the first time the implication of the microbiota in the progress of the disease [27, 28]. Moreover, despite the low resolution of the techniques employed, authors were able to characterise a particular microbial profile [29], or even particular strains [31, 32], related to CeD, different to healthy controls, and patients undergoing GFD, and the implications of the type of feeding in infants, and HLA-DQ genotype in the colonisation of some bacterial species DGGE [27, 28].

### 1.3 New high-throughput technologies

In 2012, 16S rRNA gene high throughput sequencing was used for the first time to determine the composition and temporal changes of the gut microbiota on infants at risk of CeD [33]. The main advantage of the technique is that it allows comparative analysis of the whole microbial community, diversity, and abundance at significant sequencing depths, while traditional methodologies focus only on some members of the communities [34]. The method consists of sequencing the 16S rRNA gene marker amplified from total DNA extracted directly from the tissue or environment sampled without previous culture [35]. The 16S prokaryotic ribosomal rRNA genes sequences are reliable markers for taxonomic classification and phylogenetic analysis of bacteria.

This gene is ubiquitous in the kingdom and has considerable variability in its sequence between different species, allowing their classification into different taxa. However, currently available high throughput sequencing techniques do not allow the sequencing of 16S rRNA gene amplicons spanning the gene’s full length. So, the strategy used is to sequence any of its nine hypervariable regions (V1-V9) or the regions between them [36].

On the other hand, shotgun metagenomics is based on the untargeted sequencing of genetic material present in the sample. It allows the taxonomic analysis and functional profiling of all types of microorganisms present in the microbiome (bacteria, viruses, fungi and protozoa). Moreover, it allows identifying species, strains, and even Single Nucleotide Polymorphisms (SNP) [37].

We performed a scoping review of the literature employing high-throughput sequencing of the 16S rRNA gene marker to analyse the microbiome profile in CeD. Also, we performed a merged-data analysis combining the multiple datasets of 16S rRNA gene microbiome sequencing available in public databases following the same methodology and levering proper metadata to get a consensus profiling of the microbiome in the disease. We compile datasets from different body sides and extensive metadata, considering the influence of GFD over the gut microbiome, trying to find microbial biomarkers for CeD not only in the duodenum but also in less invasive samples as saliva, stool, and oropharynx exudates.

## 2. Methods

### 2.1 Literature search, identification, and selection of relevant studies

This study follows the scoping review methodology for searching and assessment of the relevant studies [38]. This methodology adopts the following workflow (1) identification of a research question; (2) identification of relevant studies; (3) study selection; (4) charting the data; (5) collecting, summarising, and reporting the results, for this, we conducted a systematic and comprehensive literature search of the studies in years from 2010 to August 2020; that carried out metagenomic sequencing of the 16S rRNA gene for microbiomes of CeD patients, including those on a GFD as well as on an unrestricted diet. We searched on PubMed, Google Scholar, and SCOPUS. However, no additional studies to those in PubMed were found in Google Scholar or SCOPUS. The terms used for the search, the automatic filters, and the statistics results are summarised in Table S1; Figure S1 summarises the methodology used for the scoping review. The studies retrieved with the previous strategy undergo manual curation to exclude non- CeD related studies and studies using non-high-throughput 16S rRNA gene sequencing, which escaped the automatic filters.

### 2.2 Merged-data analysis of publicly available microbiome data sets

Out of the total selected studies for the literature review, those included in the merged- data analysis met the following criteria: (a) Bacterial 16S rRNA gene sequenced from total DNA using high-throughput sequencing, (b) Data must be available in fastq format in one of the publicly accessible databases NCBI Sequence Read Archive [SRA] or EBI European Nucleotide Archive [ENA]), (c) Metadata on the sequenced biological samples must be available and must include tissue of origin, information on the type of diet (with or without gluten), and clinical classification (case or control).

### 2.3 Amplicon Sequence Variant detection and taxonomic assignment

All data from the selected studies were available in the European Nucleotide Archive (ENA) database. Raw 16S rRNA gene sequencing data sets from each study were downloaded from that database. The quality of the sequencing was examined using FastQC v.11.9 software [39], and the primers used in the PCR amplification of hypervariable regions were removed using the Cutadapt v.2.9 software [40], For data obtained by paired-end sequencing, each pair of reads was joined by overlapping assembling using the software FLASH v.1.2.11 [41], Then, each study was processed individually for the construction of the ASV count table for each sample, using the software DADA2 v.1.15 [42]. Briefly, DADA2 was used for quality filtering of the sequences, detecting exact amplicon sequence variants (ASV), removing chimeras, and finally,he taxonomic assignment of ASVs with SILVA SSU v.138 database [43]. The computer code was developed with the R and “bash shell” programming environments for GNU / Linux. All the workflow and the specific criteria for each step in the analysis for each dataset are available on GitHub https://github.com/juearcilaga/Assesing-microbiome-profiles-of-celiac-disease-patients.git.

### 2.4 Data merging filtering and normalisation

As a result of the variant detection process, two tables were obtained per each study. The first one with the abundance of each variant (ASV-count table) and the second one, with the taxonomic assignment of ASVs in six taxonomy ranks (Kingdom, Phylum, Class, Order, Family, and Genus). Tables were combined using the phyloseq_merge command. With phyloseq taxa_glom command, taxa were agglomerated at the genus level, to avoid ASV duplication bias. All tables, along with raw and processed data from this study, are available in FigShare. https://figshare.com/projects/An_lisis_del_microbioma_en_Enfermedad_Cel_aca/82547

### 2.5 Statistical analysis

All the tests were carried out using the Phyloseq and Vegan v.2.5.6. [44] packages of the R programming environment. For the representation of statistical significance in the graphs, the following equivalences were used: p <’***’ -> 0.001, ’**’ -> 0.01, ’*’ -> 0.05, ’.’ 0.1, ’’ 1. The bar graphs and the scatter graphs were obtained in the R programming environment using the ggplot2 package [45].

### 2.6 Estimation of the biological diversity and composition of the microbiome

The Phyloseq v.1.3 package [46] from R v.3.6.3 [47] was used to estimate rarefaction curves for each sample. Rarefaction curves were useful to determine the minimum sequencing depth for reaching the saturation of observed species.

The diversity and richness of each sample were estimated by Shannon, Simpson, and Chao1 indexes using the phyloseq package. The Shannon and Simpson indexes were estimated after normalising the data using the Centered log-ratio method in the Compositions package [48], while the Chao1 index was estimated on the raw data. The normality of the data was evaluated using the Shapiro-Wilk test. The Wilcoxon-Mann- Whitney test was used to compare diversity means between two groups and the Kruskal-Wallis test for more than two groups. The homoscedasticity of the variances was calculated using Levene’s test. Since the Shannon index data did not meet the criteria of homoscedasticity or normality, the data were adjusted to a normal distribution by square root transformation and subsequently analysed with the Welch test.

Also, a principal coordinate analysis (PCoA) was performed on weighted Unifrac distances to display whether the data was grouped by any of the variables included in the metadata (case/control and tissue sampled, 16S rRNA gene region, age, gluten / free diet, and sequencing technology). Before PCoA data analysis, taxa with less than five counts in raw ASV-count tables or being present in just one sample were discarded.

### 2.7 Differential abundance, association, and LDA for the discovery of microbial markers on CeD

Data for each tissue was subsetted from raw ASV-count tables. The taxa present in less than 10% of samples or having less than ten counts were discarded. The CLR transformation was performed on raw ASV-count tables before differential abundance analysis with RNAseq methods and Correlation Analysis. Before the LDA Effect Size (LEfSe) analysis, taxa with less than ten counts in raw ASV-count tables were discarded. Both data rarefaction and the Total Sum Scaling (TSS) normalisation were performed on raw ASV-count tables.

The difference in taxa abundances between cases and controls for each sampled tissue was evaluated with DESseq2 software v.1.26.0 [49]. Differences with adjusted *p*-value < 0.01 and Fold change < 3 were considered significant. Association among the taxa at the genus level and CeD were studied by Spearman-Rank correlation analysis. Also, a LEfSe analysis was performed, and genera were considered significant if adjusted *p*- value <0.1 and LDA >2.

### 2.8 Inference of the microbiome metabolic potential

The software PICRUST2 [50], which infers the genes encoded in studied the taxa genomes, was used to predict the microbial communities’ functional and metabolic capacities present in each sample. The results consist of tables of genes, metabolic pathways, and enzymes abundance in each sample. Data were normalised by using the CLR transformation. The differential abundance of metabolic pathways and genes, between cases and controls, was estimated using DESeq2. Subsequently, a targeted search for differentially abundant genes whose impact on the celiac disease’s pathogenesis is of interest was carried out.

## 3. Results

### 3.1 Literature search, identification, and selection of relevant studies

We found 19 original research papers matching our criteria; we decided to group the studies into five categories according to the final aim and the main findings as follow: (1) Gut microbiome of children at-risk of CeD before the onset of the disease (2) Gut microbiome of CeD patients in comparison with healthy controls, and other groups of interest, (3) Influence of GFD on the microbiome of healthy people (gut) and CeD and Non-Celiac Gluten Sensitivity patients (saliva and gut) (4) Upper gastrointestinal tract microbiome in CeD and (5) Treatments: dietary interventions, prebiotics and hookworm infections. Table S2 summarise the main findings of the select papers.

#### 3.1.1 Gut microbiome of children at risk of CeD before the onset of the disease

We found four research in which a longitudinal study of the microbiome profile in stool samples of children at-risk of CeD before the disease’s onset was performed. Some of these studies focused on identifying microbiome composition at an early age in at-risk infants that could be predictive of CeD development [33, 51, 52], the effect of the time of first gluten exposure [33] and other environmental factors [53]; such as delivery mode, antibiotic exposure and infant feeding type on microbial gut composition and/or CeD development.

Sellitto et al. (2012) were the pioneers in using the Pyrosequencing of the 16S rRNA gene to study infant faecal microbiome before CeD onset. Their research aimed to know if gut microbiota trajectory in early life may predict the development of celiac disease and if first-time gluten exposure could affect the time of CeD onset. The authors found that the microbiota composition was highly different among at-risk infants before 18 months of age but converged at 24 months. In all children, the microbiota was characterised by a low abundance of members of the phylum *Bacteroidetes*. Infants were breastfed until six months of age. From 6 to 12 months, eight children followed a GFD and the other eight children followed a gluten-containing diet. From 12 to 24 months, all the children followed a gluten-unrestricted diet. At 12 months of age, more children in the group of early exposure to gluten presented antigliadin antibodies. Only one out of all children belonging to the group of early exposure to gluten developed CeD. The children who developed CeD showed a reduction in bacterial richness from 6 to 10 months. However, no definitive answer can be given to the original questions because only one child is insufficient to make statistical analysis and achieve valid conclusions [33].

Six years later than Sellitto et al. (2012) study was published, Rintala et al. 2018 assessed the same question. May gut microbiota trajectory in early life be a predictor of celiac disease development? But, increasing sample size and follow-up time. They studied changes in microbial communities of 27 infants genetically at risk of CeD from 9 to 12 months of age. Unlike Sellitto et al., they did not test the effect of first-time gluten exposure. The introduction of gluten was initiated from 4.4 to 5.7 months of age in all cases, similar to the early exposure time in Sellito et al. The follow-up time of this study was more extensive than in Sellito et al. Nine children (all girls) out of the 27 developed CeD at the median age of 3.5 years (range 2.6–4.2 years). These observations would not have been made in the followed time studied by Sellito et al. (0-24 months). Rintala et al. study showed a significant increase in children’s microbiota alpha diversity in general between 9 and 12 months of age. Still, no differences among healthy controls and the group who develop CeD were detected. The study concludes that the individuals who develop CeD, do not already in the early infancy, have a distinct faecal microbiota composition compared to other infants with risk-HLA-haplotype, suggesting that the onset of CeD is more likely a consequence of a solid external trigger rather than gradual development due to peculiarly vulnerable gut microbiota [52].

Another study by Olivares et al. (2018) aimed to answer the same question about the predictive power of microbial communities’ changes at an early age of infants genetically at risk of developing CeD. With a similar sample size and time of gluten exposure, samples were taken earlier in the baby’s life. They studied the microbiota of 20 children from 4 to 6 months of age. The introduction of gluten was initiated from 6 months of age onwards in all cases. Nine out of the twenty children developed CeD among 16 and 40 months of age and one at 82 months. Unlike Rintala et al. they found changes among microbiota of cases vs controls. The study showed a significant increase in microbiota alpha diversity of healthy control children (characterised by increases in *Firmicutes* families) between the ages of 4 and 6 months, which did not occur in the child group who developed CeD. An increased relative abundance of *Bifidobacterium longum* was associated with control children, while increased proportions of *Bifidobacterium breve* and *Enterococcus spp.* were associated with CeD development. Authors conclude that the diversity of microbiome increases among 4 and 6 months of age in children who remained healthy but not in those who develop CeD [51].

Finally, Leonard et al. (2020) attempted to characterise the changes in microbial communities of infants genetically at risk of CeD. However, unlike the studies mentioned above, they did not perform a follow-up, and they did not evaluate the predictive power of the microbiota composition because their question was different: Do environmental factors like delivery mode, antibiotic exposure and infant feeding type impact on the microbial composition of gut in at-risk infants?. Authors found that genetic risk to develop CeD was associated with decreased abundance of *Streptococcus* and *Coprococcus* and decreased abundance of *Veillonella*, *Parabacteroides* and *Clostridium perfringens* at 4-6 months of age. Also, with increased abundance of *Bacteroides* and *Enterococcus* species at 0 months. Cesarean section delivery was associated with a decreased abundance of *Bacteroides vulgatus* and *Bacteroides dorei* and folate biosynthesis pathway. With an increased abundance of hydroxyphenyl acetic acid, alterations implicated in immune system dysfunction and inflammatory conditions. They found differences in the microbiota regarding exposure to breastmilk and formula and between antibiotic exposure (as an environmental risk factor). Despite the study provides insights into taxonomic and functional shifts in the developing gut microbiota of infants at risk of CeD linking genetic and environmental risk factors to detrimental immunomodulatory and inflammatory effects, it is unclear whether they indeed contribute to the future development of CeD [53].

In summary, these studies evidence changes in the microbiome of infants with CeD, particularly modifications in diversity, demonstrating the importance of the inclusion of gluten in diet.

#### 3.1.2 Gut microbiome of CeD patients in comparison with healthy controls and other groups of interest

We reviewed five papers in which a case-control study of the microbiome profile in stool and/or duodenal samples of CeD patients was carried out.

In 2013, Cheng et al. performed a study analysing the duodenal microbiota from newly diagnosed children with CeD. They did not find significant differences in the abundance of taxa between groups, neither at phylum nor at the genus level. However, using Random Forest (RF) algorithm with a subpopulation of eight genus-like taxa, they were able to separate samples between CeD and healthy controls. Taxa included *Prevotella melaninoenica*, *Haemophilus* sp. and *Serratia* sp. augmented in CeD, whereas *Prevotella oralis*, *Proteus* sp., *Clostridium stercorarium*, *Ruminococcus bromii*, and *Papillibacter cinnamivorans* were augmented in controls. This classification system had an error rate of 31.6% [54] This study showed that bacteria from *Proteobacteria* were elevated in CeD and butyrate producers *Firmicutes* were diminished despite the sequencing technology used.

A different approach was conducted by Pellegrini et al. (2017). They analysed the duodenal microbiota of CeD and patients with Diabetes Type 1 (T1D), trying to find differences in the inflammatory state compared with healthy controls. The authors did not found differences in the diversity measures across the studied groups. However, they found differences in the inflammatory status of the patients studied according to the type of pathology and some relevant changes in the microbiota. In this regard, authors showed that CeD patients share some similitudes in the microbial profile with T1D *i.e.*, reduction in *Bacteroidetes* phylum and the class *Clostridia,* but the microbial profile differed according to the abundance of *Proteobacteria*, being increased in CeD, particularly *Gammaproteobacteria* from the family *Pasteurallaceae*, and the genus *Haemophilus* spp. [55].

Later, Garcia-Mazcorro et al. (2018) analysed duodenal and stool microbiota from gluten-related disorders in Mexican patients. Again, they only found differences in particular taxa abundance but not in the general parameters of diversity. Regarding duodenal samples, a lower abundance of *Bacteroidetes* and *Fusobacteria* was found in CeD patients and a higher abundance of *Actinobacillus* (*Gammaproteobacteria*), *Finegoldia* (*Clostridia*) and TM7 in Non-Celiac gluten sensitivity patients (NCGS), on the other hand, *Sphingobacterium* (*Bacteroidetes*) was higher in healthy subjects. The microbial profile of stool samples was different, *Firmicutes* were predominant regardless of disease status, and *Bacteroidetes* were less than 1%. For the first time, this study showed particular microbial changes specific to gluten-related disorders in Mexican people [56].

Bodkhe et al. (2019) studied faecal microbiome and duodenal samples from patients with CeD and first-degree relatives. Despite no significant differences in alpha diversity between sample sites were found, a different overall composition of the microbiome was inferred between groups, showing that microbiome composition and diversity were different between duodenum and stool samples for all disease status evidenced by changes in individual taxa abundance [57].

More recently, Panelli et al. (2020) studied also duodenum, stool, and saliva from CeD patients. In stool samples, *Bacteroides* were predominant, and no differences in taxa abundance were detected between groups. A decrease in *Firmicutes* and *Actinobacteria* accompanied by an increase in *Proteobacteria* was evident in the duodenum and salivary samples of CeD groups. While at the genus level an expansion of *Neisseria* and a reduction of *Streptococcus* were found in CeD groups compared to controls was also appreciated. Authors conclude that there are different patterns of diversity across groups and body tissues [58].

These studies demonstrate that alfa-diversity between samples from patients with CeD, and other groups did not show differences. However, specific changes in taxa abundance were found. Some similarities were evident across studies, for example, the increase of *Proteobacteria* in CeD patients and a decrease in *Firmicutes* and *Actinobacteria*, evidencing the existence of a dysbiosis in CeD with a predominance of gram-negative bacteria.

#### 3.1.3 Influence of GFD on the microbiome of healthy people (gut) and CeD and NCGS patients (saliva and gut)

We found five papers studying GFD in different conditions.

Bonder et al. (2016) study the influence of a GFD through time by measuring microbial changes in patients with a GFD for four weeks followed by a washout period. The authors did not observe a time-dependent change in alpha diversity or differential abundance in the microbiome composition. There was a substantial difference when comparing individuals regardless of time point, showing that inter-individual effect on the microbiome variation is stronger than the effect of diet. Carbohydrate and starch metabolisms were altered with differences between GFD and a healthy diet in the abundance of pathways related to tryptophan metabolism, butyrate metabolism, fatty acid metabolism, and seleno-compound metabolism. The authors conclude that change in diet did not influence the bacterial diversity within a sample. Inter-individual differences were more influential than the effect of GFD [59].

Garcia-Mazcorro et al. (2018) the study was partially discussed in section 1.2. They also investigated the potential microbial signatures associated with GFD by consuming certified gluten-free foods in patients with paired samples. Contrary to Bonder et al., they found that the groups studied showed group-specific variation over time after consuming the GFD for four weeks, notably NCGS presented an increase in the duodenal *Pseudomonas* on the GFD. In contrast, only half of CeD patients showed increases in *Pseudomonas*, but these increases were pronounced to affect median values. Biomarker analysis of the taxa at the genus level confirmed the results on *Pseudomonas* and showed that other *Proteobacteria* (e.g., *Stenotrophomonas* and *Novosphingobium*) were significantly more abundant on the GFD. However, these differences were only observed in duodenum samples, and there were no differences regarding these bacterial groups in faecal samples [56].

Ercolini et al. (2016) studied the influence of a change on GFD over the microbiome of celiac children following an African- style GFD during two years before the 60 days of treatment with an Italian-style gluten-free diet. After Italian-style GFD treatment, the microbiome underwent a time-dependent reduction in alpha diversity compared to baseline samples on African-style GFD. Many significant differences in microbiome composition were found. Regardless of changes in abundance, the core microbiome was conserved in children and is composed of fifteen genera belonging to the phylum *Firmicutes*. The metagenome prediction confirmed the diet’s remarkable effects at a possible functional level. At baseline, there was a higher abundance of genes related to carbohydrate metabolism. In contrast, after 60 days of diet change, the saliva was characterised by a higher abundance of genes related to amino acid metabolism, vitamins and cofactors. The change in GFD diet style alters the Saharawi children’s salivary microbiome significantly, reducing its alfa diversity, changing their beta diversity and varying the abundance of microbiome members and functional pathways [60], suggesting that diet composition is a critical factor in evaluating the effect of GFD over CeD patients microbiome.

D’Argenio. et al. (2016) compared the microbiome of CeD patients with unrestricted diet, under a GFD, and healthy controls. They found that *Actinobacteria* and *Firmicutes* phyla were less abundant in the CeD on gluten-containing diet than in the other two groups. *Betaproteobacteria* class was highly represented in active CeD patients’ gut microbiome, although its levels did not directly correlate with disease severity. *Neisseriales* order, the *Neisseriaceae* family, and the *Neisseria* genus were significantly more abundant in active CeD patients than in the other study groups. Bacterial richness did not differ between CeD and control. The Beta diversity of bacterial communities was statistically significant within the three groups, suggesting that CeD patients’ microbiomes either on GFD or unrestricted diet are more similar to each other than either of them to the control microbiomes. Authors conclude that overall microbial community composition is different among CeD and controls, although richness is comparable [61].

In a similar study using samples from duodenum, Nistal. et al. (2016) did not found significant differences in alpha or beta diversity among CeD and controls; however, *Asteuralleaceae* and *Streptococcus* members were frequent in CeD patients, mainly on a gluten-containing diet. The authors did not find dysbiosis in the microbiota composition in the duodenum. They suggested that the microbiome’s role in the disease must be associated with the function and not merely the composition [62].

In conclusion, there is no consensus in the literature regarding the adherence of CeD patients to a GFD. Some authors mention that inter-individual effect on the microbiome is greater than the effect of diet, whereas some studies report partial changes in the diversity and microbial composition after acquiring this type of diet.

#### 3.1.4 Upper gastrointestinal tract microbiome in CeD

CeD mainly affects the small intestine (duodenum), its diagnosis often comprises a combination of clinical, serological and histopathological findings, and small-bowel biopsy specimens are fundamental for an accurate diagnosis [23]. However, biopsy- sparing diagnostic guidelines have been proposed and validated in a few recent prospective studies, as the obtention of a duodenal biopsy o duodenal content is an invasive procedure, some studies aim to find microbial markers for other parts of the upper gastrointestinal tract, with minor invasive procedures (saliva. and oropharyngeal exudate) in order to identify possible microbial markers as “surrogate markers” of the duodenum. We found three papers studying the upper gastrointestinal tract microbiome in CeD.

In a study by Francavilla et al. (2014), the authors revealed that the composition of the main bacterial phyla differed between the salivary microbiota of CeD children and controls. *Lachnospiraceae, Gemellaceae* (genus *Gemella*), and *Streptococcus sanguinis* were most abundant in CeD children’s saliva. At the same time, the abundance of *Streptococcus thermophiles* was markedly decreased. Other *Firmicutes* (e.g., *Veillonella parvula*) associated with oral health were found at the highest levels in controls’ saliva. The saliva of CeD children harboured the highest levels of *Bacteroidetes* (e.g., *Porphyromonas spp*., *P. endodontalis*, and *Prevotella nanceiensis*) and the lowest levels of *Actinobacteria*. The authors conclude that the salivary microbiome of CeD children is less diverse, with a different community structure of healthy children’s microbiome. CeD microbiome has increased commensal opportunistic pathogens and reduced microbial species associated with health [63].

A similar study conducted by Tian et al. (2017) analysed salivary microbial profiles and protease activities of the microbiome given their potential involvement in gluten digestion and processing in CeD and refractory CeD (R-CeD). Significant differences were found between the CeD and R-CeD groups with respect to *Bacteroidetes* (CeD > R-CeD), *Actinobacteria* (CeD < R-CeD), and *Fusobacteria* (CeD >R-CeD) abundances. The overall microbial diversity was greater in the healthy subjects than in the CeD group, and controls had a higher abundance of TM7 sp., *Treponema sp., Simonsiella muelleri, Actinomyces sp., Porphyromonas sp Alloscardovia omnicolens* in comparison to CeD and R-CeD. The authors studied gluten degradation rates and found that gluten substrate degradation in saliva was relatively low in healthy controls. However, gluten degradation rates were significantly higher in saliva from CeD patients, regardless of whether the tripeptide substrate or the 33-mer substrates were employed [64].

Finally, Iaffaldano et al. (2018) found that the oropharynx of CeD patients showed a microbial imbalance similar to that previously described in the CeD duodenal microbiome undergoing an unrestricted diet. They observed an almost coincident oropharyngeal microbiome in controls and GFD subjects. Genes associated with polysaccharide metabolism predominated in control and CeD-GFD microbiomes. In contrast, in CeD patients with an unrestricted diet, the more significant metabolic potential for degradation of amino acids, metabolism of lipid and ketone bodies and microbial antioxidant defence mechanisms was observed. Finally, the authors suggest a continuity of CeD microbial composition from mouth to the duodenum, proposing that the characterisation of the microbiome in the oropharynx could contribute to investigating the role of dysbiosis in CeD pathogenesis [65].

Upper intestinal microbiota analysis revealed differences across CeD patients and healthy controls, suggesting the possible use of less invasive samples as a marker. There is a need to perform studies integrating a significant number of samples to determine universal markers related to CeD.

#### 3.1.5 Treatments: dietary interventions, prebiotics and hookworm infections

Due to the implications of the microbiome in CeD onset and progress, dietary modifications aiming to restore microbial eubiosis are of interest. In this sense, probiotics intake and different approaches such as hookworm infections that modify the microbiome and activate the immune response have been proposed as useful tools for modifying gut microbiota in CeD patients. We discuss here two papers evaluating both approaches.

Quagliariello et al. (2016) evaluate the probiotic effect of two *Bifidobacterium breve* strains on the gut microbiota composition of CeD under a GFD. *Firmicutes/Bacteroidetes* (F/B) ratio was calculated for each group of subjects; this ratio expresses the relationship between the two dominant phyla found in the gut microbiota and associated with several pathological conditions [66]. CeD subjects had F/B ratio values lower than the control group. However, the administration of the probiotic strains for three months showed to increase this ratio. The authors found that *Firmicutes* were significantly lower in CeD subjects not receiving the probiotic formulation than the control and probiotic groups. Similar results were found for *Actinobacteria,* underrepresented in the CeD group, and increased after the administration of probiotic strains. The treatment with the *B. breve* strains not caused changes at the levels of genus or phylum to which the probiotic belongs, but the intake of the probiotic strains acted as a “trigger” for the increase of *Firmicutes* and restoring F/B ratio. The authors concluded that three months of administration of *B. breve* strains could make the intestinal microbiota of CeD patients more similar to that of healthy individuals, restoring the abundance of some microbial communities that characterise the typical physiological condition [67].

In another study, Giacomin et al. (2016) investigated alterations in duodenal microbiota of CeD subjects before and following experimental infection with *Necator americanus and* following the administration of a low and high dose of dietary gluten. Considering that experimental infections with the human hookworm *N. americanus* could be used to treat CeD; infection alone increases anti-inflammatory cytokine, hinders colonisation of *Proteobacteria*, and favours the enrichment of *Clostridiales* in patients on GFD to improve tolerance to gluten micro-challenges [68]. They revealed that diversity in trial subjects was higher than in control subjects. Moreover, they observed significant microbial diversity changes throughout the trial: at week 24, a significant increase in diversity compared to baseline samples, whereas at week 36, diversity returned to baseline. These results suggest a correlation between infection status and exposure to escalating gluten doses and the gut microbiota composition. Finally, the authors conclude that low doses but not high doses of gluten in hookworm infected CeD patients previously in long-term GFD have beneficial effects in duodenal microbiome diversity (increase). However, the study does not differentiate between gluten and infection effects, do not measure inflammatory markers or mention symptoms and do not go in-depth on analysing individual taxa abundance changes [69].

### 3.2 Integrative data analysis of publicly available CeD microbiome data sets

The limited number of cases present in studies exploring the effects of microbiota over CeD onset, progression, symptoms development and diagnosis, impulses the need for comparison among them. However, the use of different sequencing methodologies and DNA extraction procedures difficult the comparison among results; moreover, the lack of standardisation methodologies for downstream analyses of sequencing introduces statistical biases and challenges for reproducibility and cross-study comparisons [34].

We combine 16S RNA sequencing gene datasets for the first time and perform an analysis following the same pipeline. We compared sequences generated from different regions of the 16S rRNA gene by using a reference mapping protocol for ASV assignment, in which sequences from different regions of the 16S rRNA gene will map to the same full-length reference sequence from SILVA SSU v.138 database [43] if they are from the same species. Achieving an integrative analysis including a high number of samples from different body sites and extensive metadata to find microbial biomarkers characteristic of CeD taking into account the type of diet.

We found nine out of the nineteen selected studies meeting inclusion criteria for the merged data analysis (Table 1). Table 2 shows the number of samples for each study, the clinical classification and tissue of origin (stool, duodenum, pharynx or saliva) for the data included in the analysis. Finally, we included 435 total samples, comprising 190 patients with active or treated CeD and 245 controls.

**Table 1.**
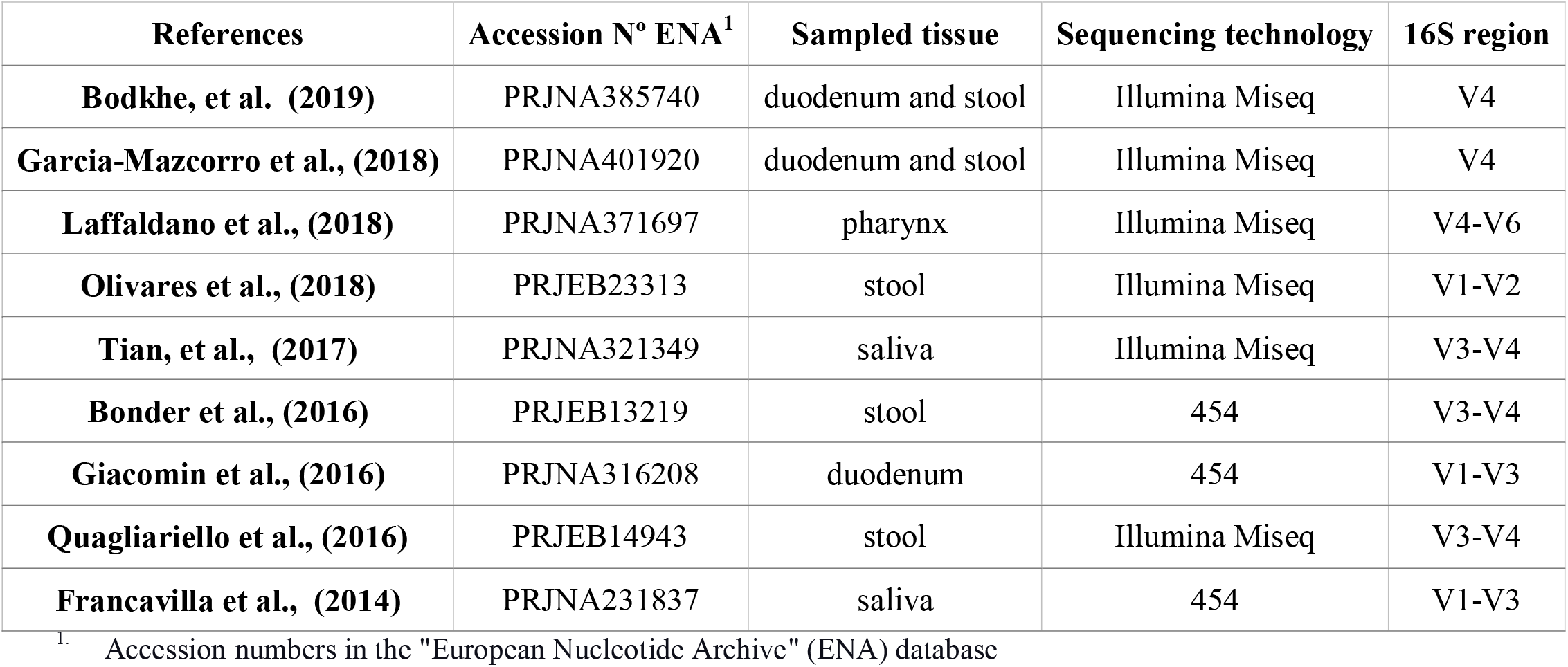
The number of samples for each study, clinical classification and tissue of origin for the data included in the analysis. The cases (n = 190) include patients with celiac disease, on a diet with or without gluten, the controls (n = 245) include non-celiac patients with gluten intolerance, gluten-free diet or without any enteropathy. The total samples add up to 435.

| References | Accession N° ENA<sup>1</sup> | Sampled tissue | Sequencing technology | 16S region |
| --- | --- | --- | --- | --- |
| **Bodkhe, et al. (2019)** | PRJNA385740 | duodenum and stool | Illumina Miseq | V4 |
| **Garcia-Mazcorro et al., (2018)** | PRJNA401920 | duodenum and stool | Illumina Miseq | V4 |
| **Laffaldano et al., (2018)** | PRJNA371697 | pharynx | Illumina Miseq | V4-V6 |
| **Olivares et al., (2018)** | PRJEB23313 | stool | Illumina Miseq | V1-V2 |
| **Tian, et al., (2017)** | PRJNA321349 | saliva | Illumina Miseq | V3-V4 |
| **Bonder et al., (2016)** | PRJEB13219 | stool | 454 | V3-V4 |
| **Giacomin et al., (2016)** | PRJNA316208 | duodenum | 454 | V1-V3 |
| **Quagliariello et al., (2016)** | PRJEB14943 | stool | Illumina Miseq | V3-V4 |
| **Francavilla et al., (2014)** | PRJNA231837 | saliva | 454 | V1-V3 |

<sup>1.</sup> Accession numbers in the "European Nucleotide Archive" (ENA) database

**Table 2.** Statistics of Input data to analyse (After filtering samples and ASVs).

| Tissue | Feeding Habit | ASVs | Samples | Cases | Controls |
| --- | --- | --- | --- | --- | --- |
| **duodenum** | GFD | 475 | 30 | 12 | 18 |
| **duodenum** | Unrestricted | 475 | 89 | 40 | 49 |
| **stool** | GFD | 374 | 89 | 52 | 37 |
| **stool** | Unrestricted | 374 | 107 | 24 | 83 |
| **saliva** | Control Unrestricted and Case GFD | 120 | 78 | 40 | 38 |
| **pharynx** | Control Unrestricted and Case GFD | 71 | 42 | 22 | 20 |

ASV= Amplicon Sequence Variant. GFD= Gluten free diet. Cases=Samples from patients with Celiac disease.

**Table 3.** Statistics of significant results obtained on Picrust2 predictions analysis.

| Tissue | Feeding Habit | Gene Differential abundance | Gene Correlation | Pathway Differential abundance | Pathway Correlation |
| --- | --- | --- | --- | --- | --- |
| duodenum | GFD | 412 | 0 | 13 | 0 |
| duodenum | Unrestricted | 404 | 0 | 8 | 0 |
| stool | GFD | 31 | 69 | 0 | 1 |
| stool | Unrestricted | 261 | 0 | 2 | 0 |

GFD= Gluten-free diet. FC>=3, *p* <0.01

#### 3.2.1 Diversity and microbial composition

Similar to previously discussed studies in section 2, we did not find differences among healthy controls and CeD patients regarding alpha diversity indexes. However, when we consider GFD (for duodenum and stool samples), we found an increment in the diversity in patients as in healthy controls undergoing a GFD. Alpha diversity of the microbiome was estimated using the Chao1, Shannon and Simpson indices (Figure 2).

**Figure 2.**
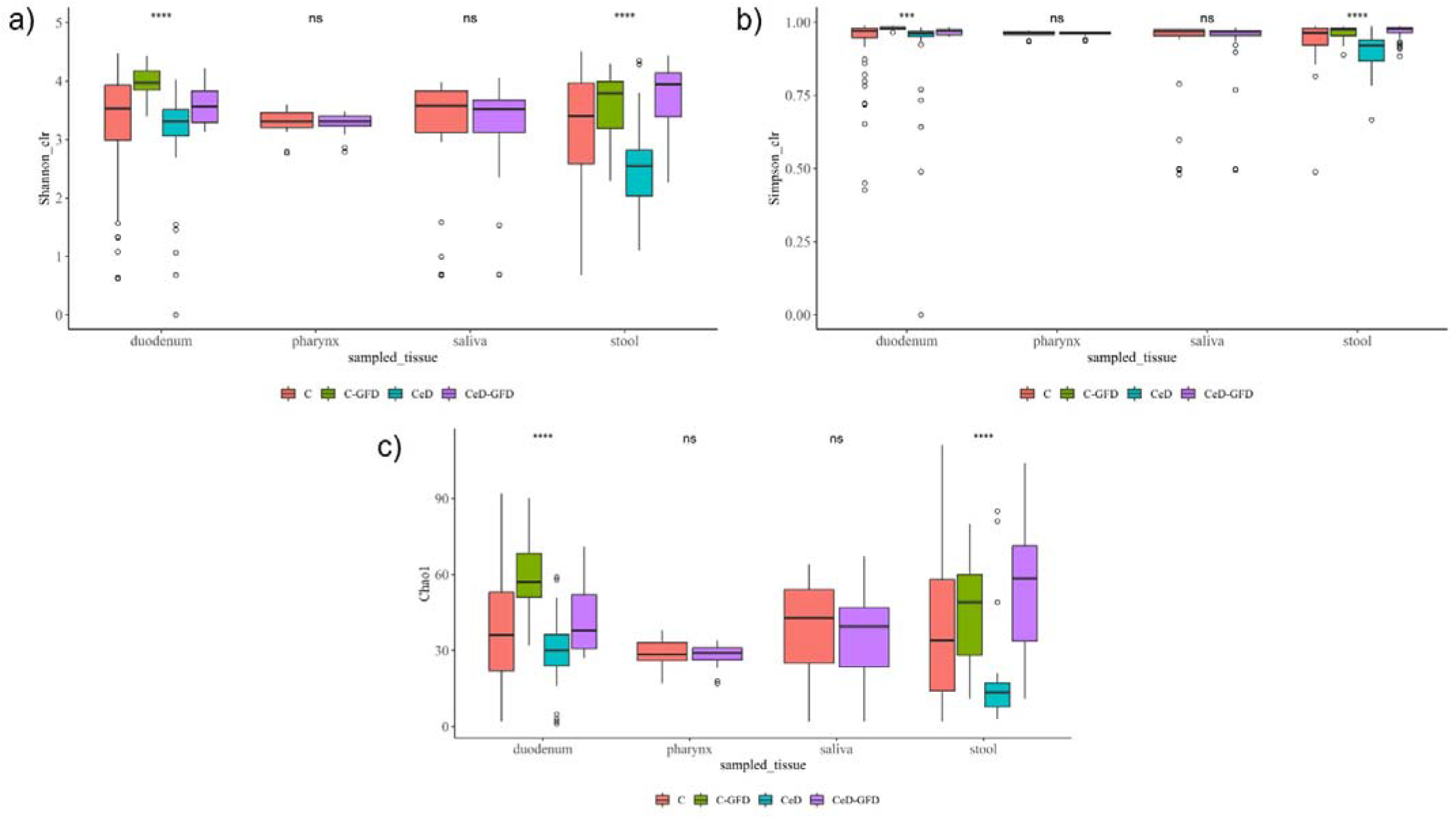
Shannon (A), Simpson (B) and Chao1 (C) diversity indices for (1) the pooled samples in cases and controls for each sampled tissue according to diet. The p-value was calculated using the Wilcoxon test in A and B and the Welch’s t-test in C. The limits of the rectangle indicate the 25th and 75th percentiles and the horizontal bars indicate the median, in C, the median equals average. Vertical bars indicate upper and lower distribution limits, and dots represent mild outliers. See in methods the codes of significance of the p-value.

For the beta-diversity analysis, we study differences among groups in each tissue sampled by PCoA using weighted UniFrac distance, but we did not find a clear separation when analysing biological variables (age, type of diet or clinical condition), indicating that there are no differences in beta diversity between the microbial communities studied and that the differences in the microbiome are not due to global differences in the abundance of the phylogenetic groups present, but probably due to the abundance of specific taxa.

Our results are in the same line as others that assure that differences in the microbiota across different body sites are enough for grouping samples [72]. Permutational multivariate analysis of variance (PERMANOVA) for each variable by ADONIS function revealed that Study Accession, in the case of the duodenum, saliva, and stool, was a factor influencing the grouping of samples. This may be explained by the experimental protocols used in each study, including differences in the sequencing platform, the region of 16S rRNA targeted, and the DNA extraction technique used, suggesting that some particular protocols may induce some biases, however, the principal coordinates explain a low percentage of the variance between the samples, indicating that other non-technical factors are mainly responsible for the variance.

#### 3.2.2 Differential analysis of the microbiota, correlation and biomarker finding

Celiac disease (CeD) mainly affects the small intestine (duodenum); according to the European Society for the Study of Coeliac Disease (ESsCD) guidelines, the diagnosis of CeD relies on a combination of clinical, serological and histopathological findings and small-bowel biopsy specimens are fundamental for an accurate diagnosis [23]. In this regard, we study duodenal microbiota intending to find microbial markers associated with this disease. On the other hand, biopsy-sparing diagnostic guidelines have been proposed and validated in a few recent prospective studies, as the obtention of a duodenal biopsy o duodenal content is an invasive procedure, we aim to find microbial markers for other parts of the gastrointestinal tract, with less invasive procedures (saliva. stool, oropharyngeal exudate) in order to identify possible microbial markers as “surrogate markers” of the duodenum.

To establish microbial biomarkers, first, we conduct a biomarker finding analysis using the LEfSe tool, followed by a differential abundance analysis using DESeq2 to identify ASVs that were differentially expressed according to studied groups. Finally, a correlation analysis looking for an association between CeD and microbial composition was performed. Figure 3 summarise the main findings obtained in each tissue analysed.

**Figure 3:**
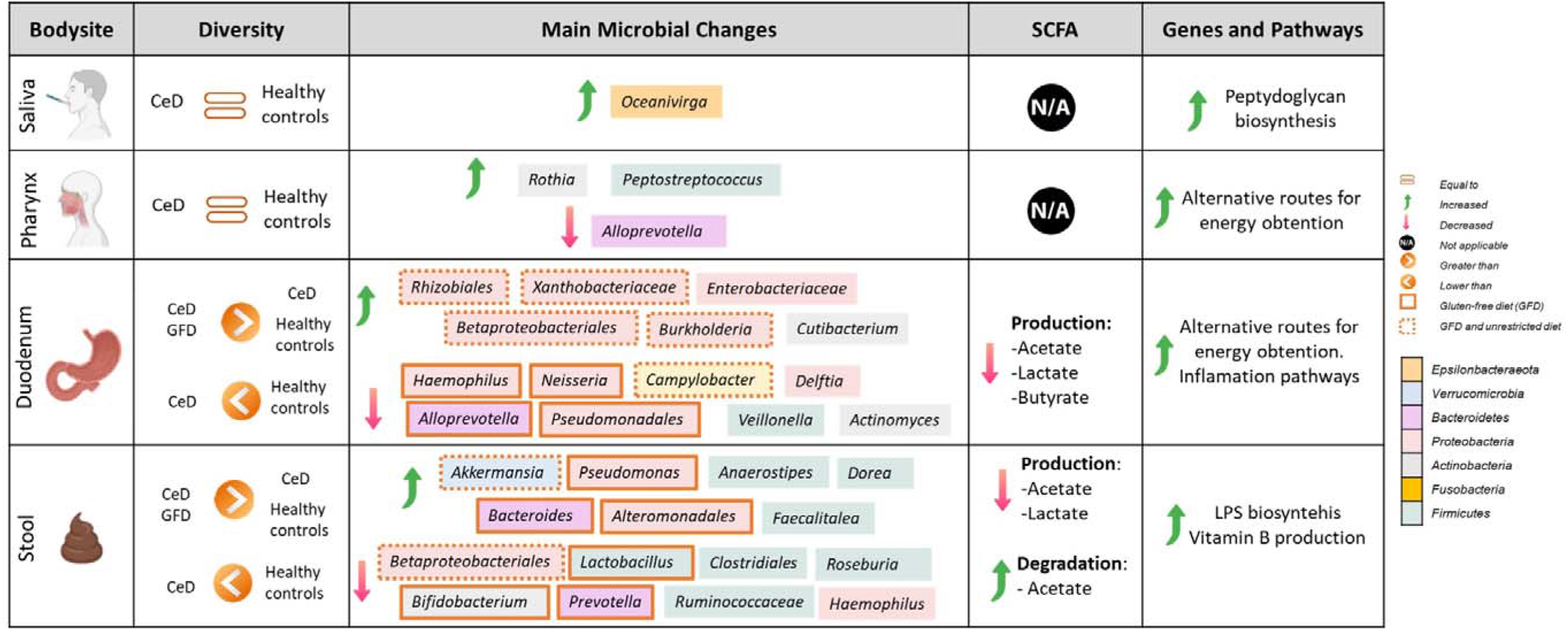
Summary of microbial and functional changes observed after data integration. SCFA= short-chain fatty acids. CeD=Celiac disease patients.

##### 3.2.2.1 Microbial changes associated with duodenal microbiota in CeD

We found that bacteria of the phylum *Proteobacteria* were characteristic of CeD patients duodenum, with different genus present according to the type of diet.

**(i) Microbial changes of duodenal microbiota from untreated CeD patients undergoing an unrestricted diet.** For duodenal samples of patients undergoing an unrestricted diet, we found 23 ASV with an LDA score greater than 3 (Figure 4a). Nine ASV were associated with CeD patients mainly from *Proteobacteria* phylum, particularly bacteria from the *Burkholderia-Paraburkholderia-Caballeronia* clade, *Alphaproteobacteria*, and *Enterobacteria*, and *Actinobacteria* from the family *Corynebacteriaceae*. On the other hand, healthy controls were enriched in *Firmicutes* phylum class *Negativicutes*, and phylum *Epsilonbacteraeota*.

**Figure 4:**
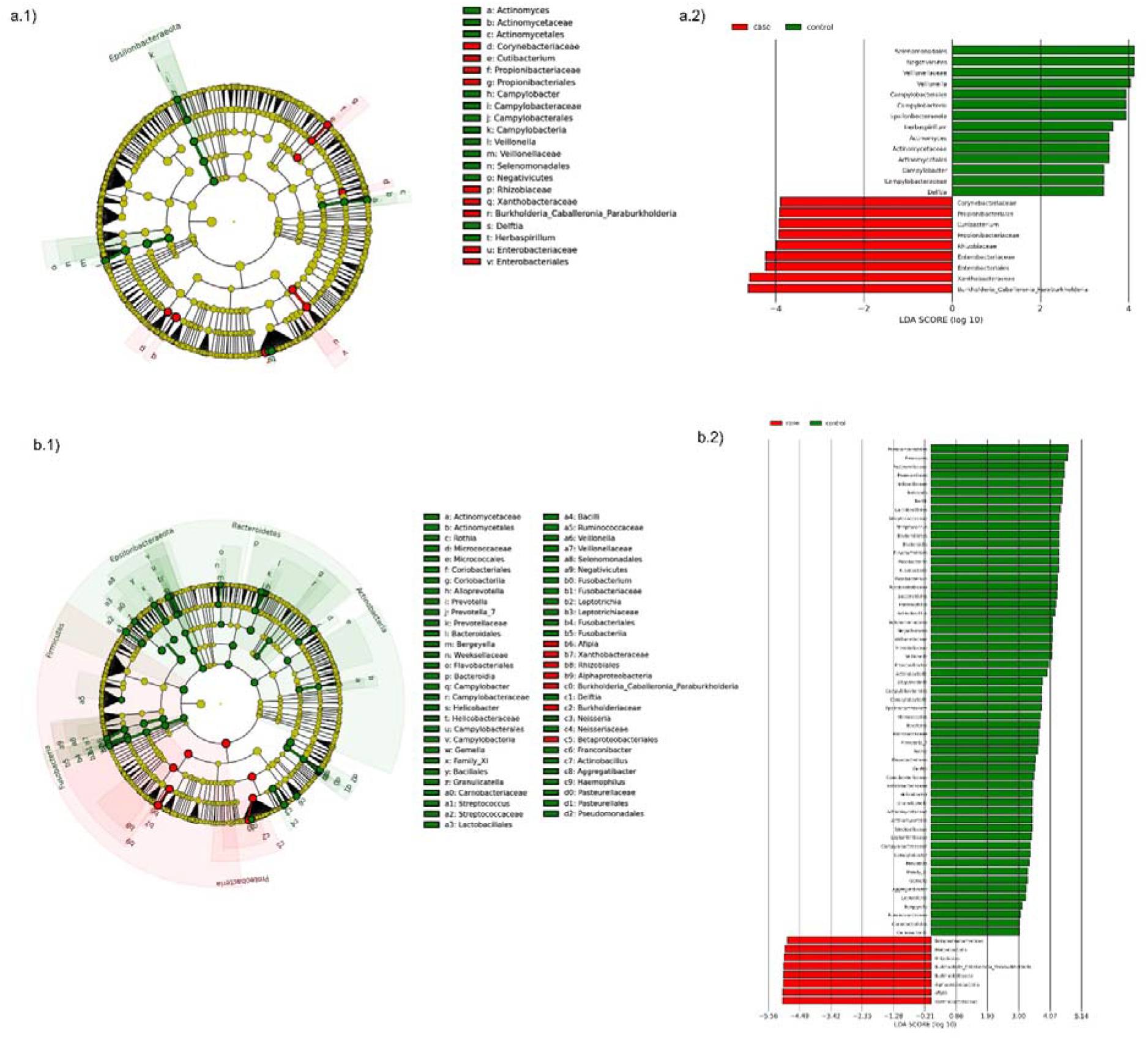
Linear discriminant analysis (LDA) integrated with effect size (LEfSe). Cladogram representing the differentially abundant taxonomic groups in the microbiota from the duodenum of CeD following an unrestricted diet (a.1) or a GFD (b.1). Graph bar representing the microbial biomarkers found in the microbiota from the duodenum of CeD following an unrestricted diet (a.2) or CeD patients following a GFD (b.2) (LDA score >3, *P* <0.001). Control: Control group, represented in green. Case: CeD patients, represented in red.

After differential relative abundance analysis, we found 93 ASVs with significant changes in abundance (FDR <0.01) (Figure 5a). Among them, 61 were decreased in CeD, and 11 were increased in CeD. Significant ASVs belong to phyla: *Firmicutes, Bacteroidetes, Proteobacteria, Epsilonbacteraeota, Actinobacteria, Spirochaetes, Fusobacteria, Synergistetes*.

**Figure 5.**
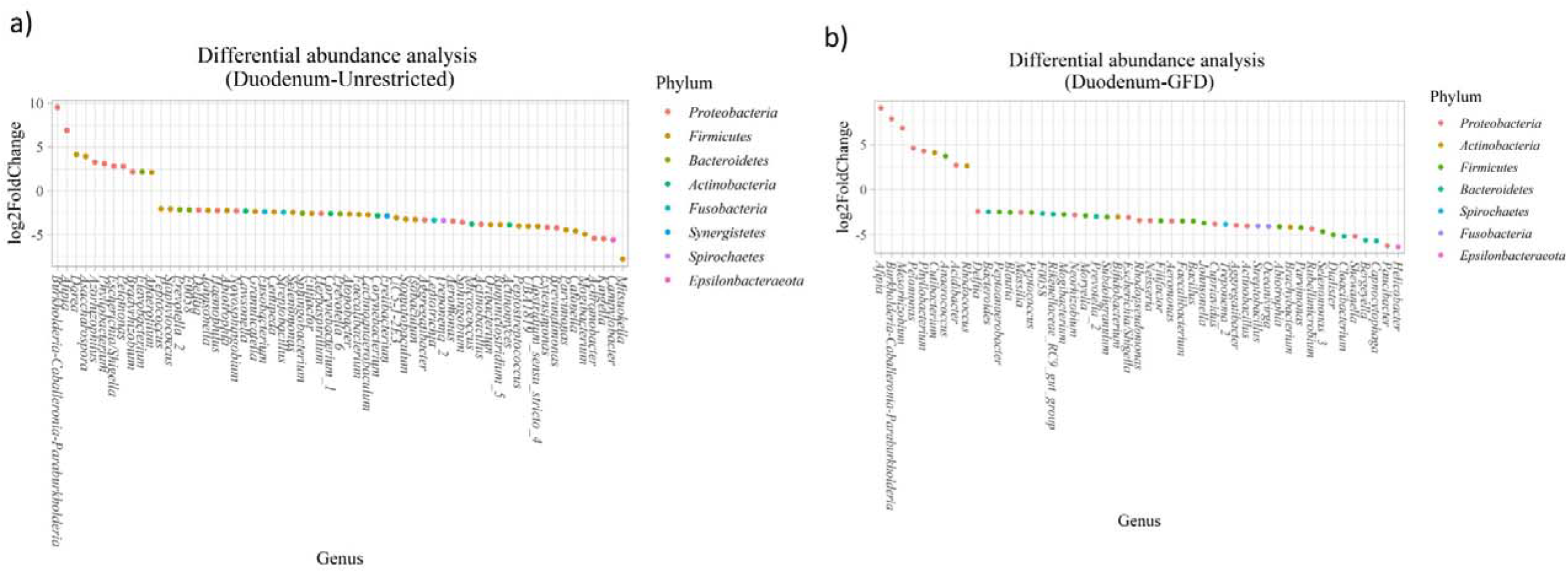
Differential abundance analysis performed on samples from the duodenum of CeD patients following an unrestricted diet (a) and CeD patients under a GFD (b) compared with healthy controls.

Finally, we found a negative association between CeD and the genus *Actinomyces,* whereas the top five positive associated genus were *Coprococcus* 3, *Hydrocarboniphaga*, *Ruminococcaceae* UCG_010, *Cutibacterium*, and *Deinococcus*.

**(ii) Microbial changes of duodenal microbiota from CeD patients undergoing a GFD.** We select healthy controls and patients undergoing a GFD to study the microbial composition between the two groups. Biomarker finding analysis revealed 65 ASVs with a LDA score greater than 3 (Figure 6a). Eight ASV were associated with CeD patients, mainly from *Proteobacteria* phylum belonging to *Burkholderia- Paraburkholderia-Caballeronia* clade, and alfaproteobacterias *Afipia*, and order *Rhizobiales*. On the other hand, 57 ASV were related with healthy controls following a GFD, particularly from phylum *Firmicutes*, *Fusobacterium*, *Actinobacteria*, and *Epsilonbacteraeota* comprising the genus *Leptotrichia*, *Fusobacterium*, *Rothia*, and *Campylobacter*, and other *Proteobacteria* (*Neisseriaceae*, *Pseudomonales* and *Haemophilus*).

**Figure 6:**
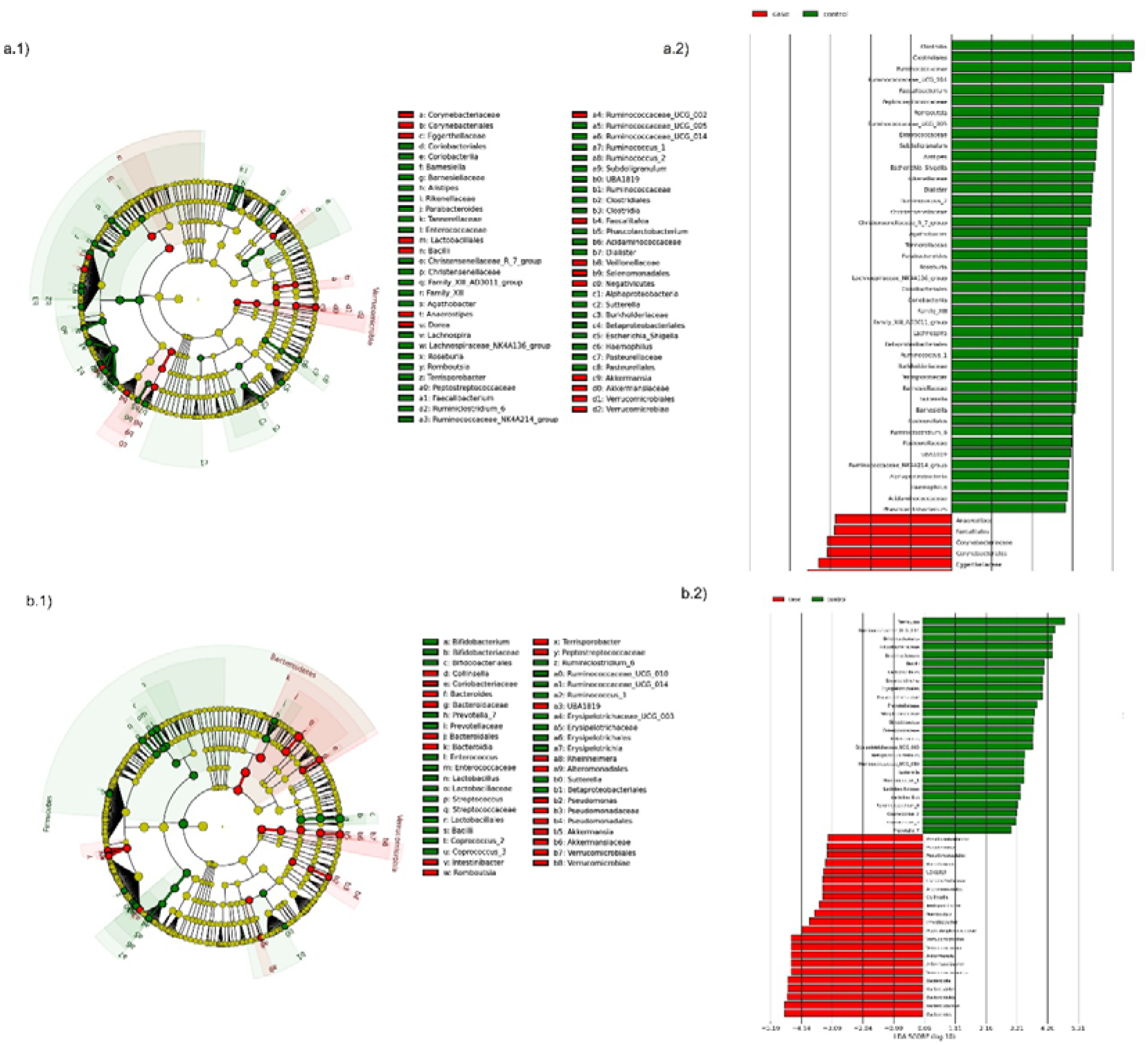
Linear discriminant analysis (LDA) integrated with effect size (LEfSe). Cladogram representing the differentially abundant taxonomic groups in the microbiota from the stool of CeD patients following an unrestricted diet (a.1) or a GFD (b.1). Graph bar representing the microbial biomarkers found in the microbiota from the stool of CeD patients following an unrestricted diet (a.2) or CeD patients following a GFD (b.2) (LDA score >3, *P* <0.001).

After performing the differential abundance analysis, we found 49 ASVs with significant changes in abundance (FDR <0.01) (Figure 5b). Nine were increased in CeD, whereas the other 40 ASV were decreased. Significant ASVs belong to phyla: *Actinobacteria, Proteobacteria, Firmicutes, Epsilonbacteraeota, Bacteroidetes, Fusobacteria,* and *Spirochaetes*.

Finally, the top five genus negative associated with CeD duodenum of patients undergoing GFD were *Haemophilus*, *Neisseria*, *Alloprevotella*, *Fusobacterium* and *Delftia*. In contrast, the top five positive associated genus were mainly from *Proteobacteria*, *Alfaproteobacteriales: Falsirhodobacter*, *Asinibacterium*, *Azonexus*, and *Blastomonas*.

Higher rates of *Proteobacteria* are linked to dysbiosis and have been suggested to increase low-grade inflammation, affecting tight-junction and leading to leaky gut [70]. This phylum was overrepresented in duodenal samples from CeD patients particularly the *Burkholderia-Paraburkholderia-Caballeronia* clade. It is worth mentioning that *Enterobacteriaceae* was overrepresented in the duodenum of untreated patients, but were not observed in patients undergoing a GFD, in fact some genus previously associated with active CeD i.e *Haemophilus*, *Neisseria*, and bacteria from the class *Pseudomonadales* was also disminished in patients undergoing GFD. These changes suggest beneficial variations in the microbial composition of the duodenum after adherence to GFD. However, these changes were not able to fully restore the microbiota, evidenced by the decrease of SCFA producers such as *Ruminococcaceae*, and *Prevotella*, and by the overrepresentation of *Betaproteobacteriales*.

##### 3.2.2.3 Microbial changes associated with stool microbiota in CeD

Stool samples are more representative of the colonic microbiota; however, they could also indicate changes related to the disease. Like in duodenum samples, we found an increase in bacteria of the phylum *Proteobacteria*, but different genus was enriched according to the type of diet. Specific changes found were as follows:

**(i) Microbial changes of stool microbiota from untreated CeD patients undergoing an unrestricted diet.** After biomarker analysis, we found 60 ASVs with an LDA score greater than 3 (Figure 6a). Of them, 17 were associated with CeD, bacteria from phylum *Verrucomicrobia* and *Firmicutes* were characteristics of this group, mainly the genus *Akkermansia*, *Anaerostipes*, *Faecalibacteria* and *Dorea*, on the other hand in healthy controls undergoing an unrestricted diet ASV from phylum *Proteobacteria* (*Betaproteobacteriales*) and *Firmicutes* mainly order *Clostridiales* were overrepresented.

Differential abundance analysis revealed 74 ASVs belonging to phyla: *Firmicutes, Bacteroidetes, Actinobacteria, Proteobacteria* with significant changes in abundance (FDR <0.01) (Figure 7a). Among them, 56 were decreased in CeD and seven were increased in CeD. Finally, top five genus positively associated with CeD were: *Achromobacter*, *Flavisolibacter*, *Geodermatophilus*, *Candidatus Rubidus*, and *Tepidimonas*.

**Figure 7:**
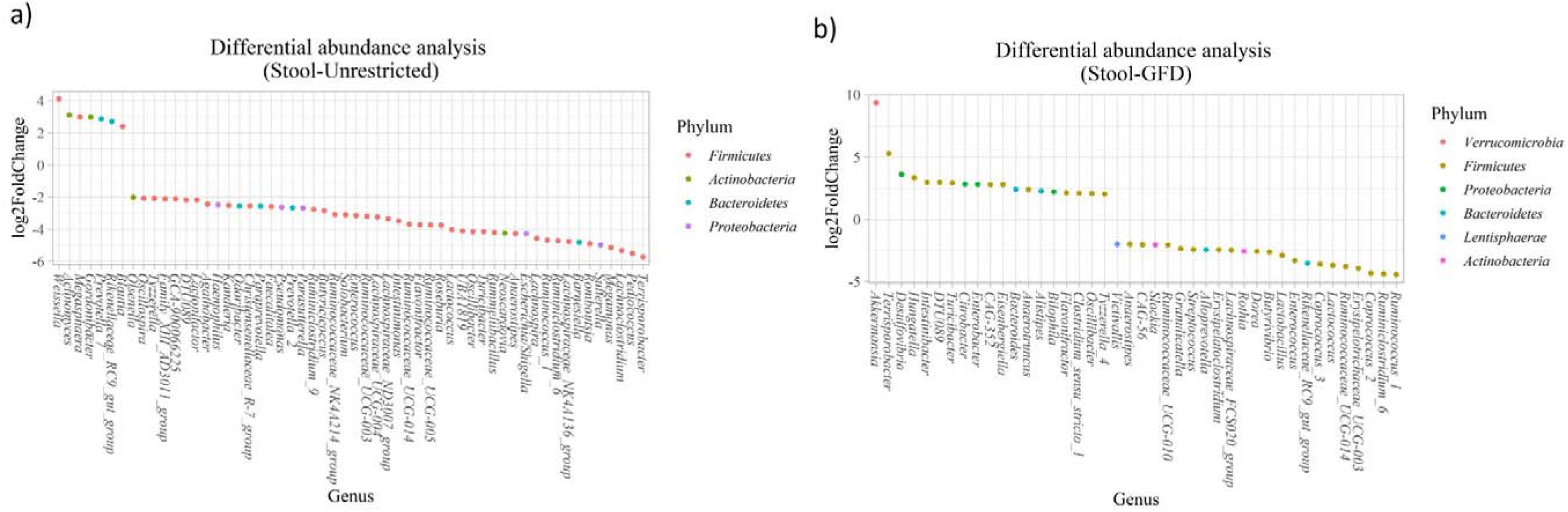
Differential abundance analysis performed on samples from the stool of CeD patients following an unrestricted diet (a) and CeD patients under a GFD (b) compared with healthy controls.

**(ii) Microbial changes of stool microbiota from CeD patients undergoing a GFD.** Biomarker finding analysis revealed the presence of 48 ASVs with LDA > 3 (Figure 6b). Twenty-two were associated with CeD patients, mainly from phylum *Verrucomicrobia, Bacteroides, Firmicutes* and *Proteobacteria*, particularly genus *Akkermansia*, *Bacteroides, Romboutsia*, and *Pseudomonas*. On the other hand, 26 ASV were related with controls undergoing GFD, mainly bacteria from phylum *Firmicutes* and *Actinobacteria* from genus *Lactobacillus*, *Streptococcus*, *Ruminoccocous* and *Bifidobacterium*. Differential abundance analysis revealed changes in 77 ASVs (FDR <0.01) (Figure 7b). Among them, 41 were decreased in CeD and 19 were increased in CeD. Significant ASVs belong to phyla: *Firmicutes, Proteobacteria, Bacteroidetes, Verrucomicrobia, Actinobacteria, Lentisphaerae*.

Finally, correlation analysis showed that 15 ASV with a significant correlation with CeD. 5 have a negative correlation with CeD (*Ruminococcus* 1*, Ruminococcaceae* UCG_014*, Ruminiclostridium* 6*, Coprococcus* 2*, Enterococcus*) and ten ASV have a positive correlation, and the top five genus positive correlated were *Intestinibacter*, *Akkermansia*, *UBA1819*, *Flavonifractor*, and *Terrisporobacter*.

It is worth mentioning that *Betaproteobacteriales* from family *Burkholderiacae* were overrepresented in the duodenum of CeD patients. However, we found that these bacteria reduce its abundance in stool and diminished notably after treatment with GFD. Also, we found an enrichment in *Akkermansia*, a bacteria associated with anti-inflammatory properties and metabolic health [71]. Despite the beneficial changes observed, stool microbiota of patients undergoing GFD presented a decrease in *Lactobacillus,* and *Bifidobacterium*, SCFA producers bacteria, which also digests intact gluten proteins reducing cytotoxicity and inflammatory responses [15, 72], and the presence of bacteria associated to CeD such as *Bacteroides*, and *Pseudomonas* both genus with species containing metalloproteases able to partially digest gluten, increasing the immunogenic effect of gliadins inducing more severe inflammation [73, 74].

We found some similitudes in stool microbiota compared to the duodenum, first an increase in *Proteobacteria*, the presence of bacteria associated with CeD (*Bacteroides*, *Neisseria*, or *Pseudomonas*) and a decrease in beneficial SCFA-producers bacteria. These findings suggest that the lower gastrointestinal tract microbiota from CeD patients is enriched in *Proteobacteria* with potential implications in activation of inflammation and immune response and a decrease in beneficial SCFA producers with anti-inflammatory potential, suggesting that gut microbiota favours the inflammatory state of the disease.

##### 3.2.2.4 Microbial changes associated with pharynx microbiota in CeD

Anatomically, the pharynx is part of the upper gastrointestinal tract, directly connected with the oesophagus, and is conventionally divided into the nasopharynx, oropharynx, and hypopharynx. Usually, oropharynx exudate, a sample comprising the part of the throat at the back of the mouth behind the oral cavity, are used for clinical diagnosis of microbial infections [75], being a valuable method for studying microbiota from the upper intestinal tract with a less invasive procedure than a biopsy [76, 77].

We found two microbial biomarkers characteristic of the pharynx from patients with CeD. *Rothia*, a nitrate-reducing bacteria usually found in the oral cavity of humans [78], and *Peptosptreptoccus*, an oral pathogen recently associated with Colorectal Cancer development [79] (Figure 8a). On the other hand, differential abundance analysis showed significant changes in abundance (FDR <0.01) of five ASV. *Veillonella, Mogibacterium (p_Firmicutes), Streptobacillus (p_Fusobacteria)* and *Mannheimia (p_Proteobacteria)* where increased in CeD whereas *Treponema 2* (p_ *Spirochaetes*) was decreased.

**Figure 8:**
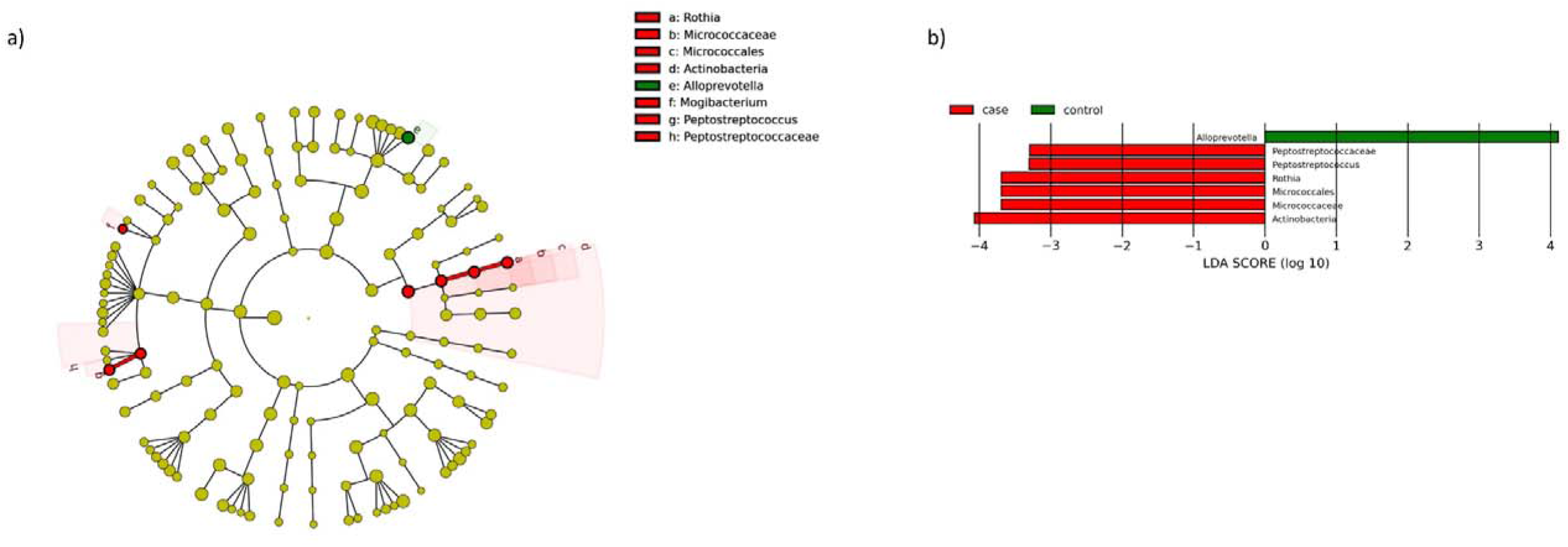
Linear discriminant analysis (LDA) integrated with effect size (LEfSe). a) Cladogram representing the differentially abundant taxonomic groups in the microbiota from the pharynx of CeD patients. b) Graph bar representing the microbial biomarkers found in the microbiota from the pharynx of CeD patients (LDA score >3, *P* <0.001).

These changes revealed a particular microbial composition from the pharynx of CeD patients, mainly associated with proinflammatory bacteria (i.e. *Mogibacterium*) and opportunistic pathogens (i.e. *Peptostreptococcus, Streptobacillus*). Although some *Rothia* species have gluten-degrading enzymes [14], the role of this bacteria in CeD pathogenesis remains unexplored. In our analysis, we found it on non-symptomatic patients undergoing GFD. Further studies will be helpful to discriminate its role in CeD and its possible use for monitoring the disease.

##### 3.2.2.5 Microbial changes associated with microbiota from saliva samples in CeD

The use of saliva samples would be beneficial for diagnosis and monitoring the disease, mainly in children, due to the non-invasiveness of the sample collection procedure. Despite the most significant changes found in the microbial composition on other parts of the gastrointestinal tract, we only found one genus related to CeD in saliva and was identified after differential abundance analysis and biomarker finding, namely genus *Oceanivirga* belonging to the phylum *Fusobacteria*, family *Leptotrichiacea*.

*Leptotrichiaceae* are relatively poor studied gram-negative bacteria, facultative or obligate anaerobes, found as colonisers of mucous membranes in the oral cavity of humans and other animals [80]. Particularly, genus *Oceanivirga* has been isolated in subgingival samples from patients with periodontitis [81]. We found this genus as a marker of CeD patients for the first time; validation as a microbial marker of CeD need to be performed.

### 3.3 Prediction of the metabolic functions profiles in bacterial communities

The functional potential of the microbiome in the different tissues was predicted using the PICRUSt tool, followed by a differential abundance analysis performed with Deseq2 and correlation analysis.

#### 3.3.1 Prediction of short-chain fatty acid (SCFA) production genes and inflammation-related pathways

To study the metabolic potential of the microbiome, the analysis was first carried out for all the genes and pathways detected and, since the intestinal microbiome, after its contribution to the digestive process of food, produces short-chain fatty acids that influence the maturation, maintenance, and behaviour of the mucosal immune system, we focus on the genes and pathways involved in the production of these compounds. Figure 3 summarises the main findings observed according to each tissue analysed.

First, we search for the 26 genes reported as potentially involved in the biosynthesis pathways of the main SCFAs produced by the human microbiome (butyrate, acetate, and lactate), given their involvement in the modulation of the immune system. Genes included were those identified by Zhao et al. (2019) by gene deletion and overexpression experiments in *E. coli*. These authors identified six genes needed in acetate production (*menI, tesA, yciA, fadM, tesB, ybgC*), eight genes used in butyrate (*entH, tesA, ybgC, ybhC, yicA, menI, yigI* y *tesB*) production, and two genes required for lactate (*mgsA* y *lldD*) production in addition to the previously known genes *pta-ackA*, *ptb-buk, ldhA, poxB, eutD, tdcD, dld* y *ykgF* [82]. Table 2 shows the number of genes found according to each tissue and the differentially abundant genes/routes between CeD patients and healthy controls.

Differential abundance analysis of metabolic pathways is summarised in Table S3, it revealed an increase in the duodenum of CeD patients, of the degradation of D-glucarate, L-arabinose, D-galactarate and biogenic amines and a decrease in the degradation of lactose and galactose. Since the cells involved in the immune response require a large amount of energy for their activation, proliferation and recruitment of other cells during inflammation, the availability of carbohydrates is scarce for bacteria to synthesise their components. Therefore, they obtain energy by alternative routes, for example, from acetate through the glyoxalate cycle, or obtaining energy are compounds such as D-glucarate, D-galactarate and biogenic amines [83]. Regarding genes involved in the production of SCFA, we found a decrease in the abundance of genes involved in the production of acetate (*ackA*) and lactate (*ldhA*) and a negative correlation with genes involved in the production of acetate (*pta*), lactate (*lldD, ldh, dld* and *tesB*) and butyrate (*tesB*). A decrease in the abundance of the fermentation pathway from hexitol to SCFAs involving the *ackA* and *pta* genes was consistently found.

In stool samples of patients with CeD, the synthesis routes of lipopolysaccharide (LPS) components of the membrane of gram-negative bacteria, acetate degradation routes, and vitamin B synthesis (B1 and B9) were found to be increased, which is consistent with our findings in the taxonomic analysis revealing an increase in the abundance of *Proteobacteria*. LPS are key factors in the activation of the immune response through TLR4 signalling, contributing to inflammation and loss of intestinal permeability seen in CeD [1, 70]. Regarding genes implied in SCFA synthesis, correlation analysis revealed that two genes related to acetate and lactate production (*pta* and *dld*) were negatively associated with CeD (FDR ≤0.01; correlation ≥0.2).

We found a generalised low presence of SCFA in microbiota of duodenum and stool from patients with CeD. The reduction of these compounds is associated with a reduction in bacterial genus that produces them, such as *Ruminoccous*, *Veillonella*, and *Clostridiales* (as observed in taxonomic analysis) and potentially with a lower fibre intake typical of a gluten-free diet [25]. Low abundance of SCFA has been previously reported in CeD at an early age [84], and may be involved in the predisposition to the development of CeD [85].

No statistically significant differences were found in the abundance of any of the 26 genes related to SCFA production regarding saliva and pharynx samples. However, in saliva samples of CeD patients, peptidoglycan synthesis was increased. Peptidoglycans are part of the cellular wall of bacteria and, besides LPS, are responsible for microbial- related inflammation mediated by innate immune response [86]. On the other hand, we found an increase in the intermediate degradation routes of aromatic compounds and amino acids in the pharynx samples, both routes associated with energy acquisition from bacteria [83].

#### 3.3.2 Prediction of genes and genes coding for prolyl peptidase enzymes

Some components of the microbiota can express enzymes different from those produced by humans and promote the digestion of compounds such as the immunogenic peptides of gluten [14, 15]. We analysed four enzymes involved in the degradation of immunogenic gluten peptides that could be involved in reducing CeD symptoms (general N-type aminopeptidase (PepN), X-prolyl dipeptidyl aminopeptidase (PepX), endopeptidase (PepO) and endoprolyl peptidase (PREP)). However, none of the four enzymes involved in the degradation of gliadin peptides were found to be differentially abundant between cases and controls or associated with CeD in any of the tissues.

## 4. Conclusions and future perspectives

Our study provides an unprecedented metanalysis of metataxonomic data from CeD patients, including (n=435) samples from CeD patients and healthy controls for different body sites. One of the most critical limitations currently found in literature is the lack of standardised methodologies for downstream analyses of sequencing approaches, introducing statistical biases and subsequent challenges for reproducibility and cross-study comparisons [34]. Despite some attempts to standardise methods, a gold standard of microbiome research is not established [87]. Herein, we create an extensive dataset processed following the same methodology and levering metadata to solve this issue; moreover, we made data publicly available (https://figshare.com/projects/An_lisis_del_microbioma_en_Enfermedad_Cel_aca/82547).

We were able to find microbial markers related to CeD at different body sites, including SCFA production and the prevalence of inflammatory pathways (Figure 3). Our results showed coordinated changes throughout the gastrointestinal tract, with specific changes according to each body site. In the upper gastrointestinal tract, we found an enrichment in *Actinobacteria* from genus *Rothia* in pharynx, and *Cutibacterium* in duodenum, and a marked decrease in *Alloprevotella* spp. in both sites. Whereas the lower gastrointestinal tract presents more changes characterised by an increase in *Proteobacteria* and a decrease in *Actinobacteria*, *Campylobacter* and SCFA producers such as *Ruminococaceae*, and *Clostridiales*. Moreover, we define some differences in gut microbiota (from duodenum and stool samples) of untreated CeD patients following an unrestricted diet vs CeD patients following GFD. Although we could not find restoration in microbial dysbiosis, we found that patients following a GFD have an overall higher microbial diversity and an abundance of certain bacteria usually related to health benefits such as *Akkermasia* [71], and a decrease in the presence on *Enterobacteriaceae*. This finding is interesting because differences in patients following GFD may be observed. Recent studies demonstrate that long-term diet is the primary driver of gut microbiota composition [65], so it is expected that following a GFD characterised by changes in protein, fat, and carbohydrate consumption [24] will have an impact on the microbial communities composition. In fact, adherence to a GFD leads to the healing of the duodenal mucosa and the resolution of symptoms and signs of malabsorption CeD [23]. However, up to 30% of patients with CeD may show signs, symptoms or persistent enteropathy after one year on a GFD, having the so-called non-responsive CeD [26] Non-responsive CeD has been poorly studied, and still, there is no clear cause-effect association to explain this entity; microbiota may play an essential role in its pathogenesis.

Previous studies have shown that *Proteobacteria* was enriched in CeD [54–58], we recognise the same finding, but we also were able to discern between differences in the abundance of particular *Proteobacteria* families according to the type of diet of the patients. For untreated CeD patients, *Burkholderiaceae*, *Xantobacteriaceae*, and *Enterobacteriaceae* were enriched, whereas in patients under a GFD we found a decrease in some *Enterobacteriaceae* (duodenum), and *Betaproteobacteriales* (stool). *Proteobacteria* may be acting as enhancers of the immune response in CeD due to the activation of the innate immune system via TLR receptor mediated by LPS [2, 8, 20].

Future prospective studies will provide the “solution” to who was before the chicken or the egg? i.e., Dysbiosis led the disease or the disease produce dysbiosis?, Current studies have a cross-sectional design and perform descriptive association at a snapshot of time, CeD is the perfect scenario for studying implications of microbiota over the pathophysiology of a disease, the trigger of the pathology (*i.e*: gluten) is traceable, and the genetic environment predisposing to the disease is also knowledgeable.

Finally, there is a need to generate larger datasets and properly apply machine learning (ML) that would help to generate helpful and more universally applicable results [88]. In this sense, future studies applying ML would be beneficial to corroborate the microbial and functional markers found in this study, and know the role in the prevalence of symptoms associated with CeD. Moreover, evaluating the usefulness of stool, oropharyngeal exudates, and saliva as surrogate markers of the microbial state of the duodenum for the diagnosis and management of CeD patients would be necessary. Finally, the potential efficacy of gut microbiota modulators such as probiotics and prebiotics as adjuvant treatment for CeD need to be further assessed.

## Availability of data and material

The datasets generated during and/or analysed during the current study are available available in FigShare. https://figshare.com/projects/An_lisis_del_microbioma_en_Enfermedad_Cel_aca/82547

The computer code was developed with the R and “bash shell” programming environments for GNU / Linux, The workflow and the specific criteria for each step in the analysis for each dataset are available on GitHub https://github.com/juearcilaga/Assesing-microbiome-profiles-of-celiac-disease-patients.git

## List of abbreviations

CeD: Celiac disease
GFD: gluten-free diet
TG2: transglutaminase type 2
APC: antigen presenter molecules
ENA: the European Nucleotide Archive
ASV: exact amplicon sequence variants
PCoA: principal coordinate analysis
CLR: Centered log ratio Normalization
LEfSe: LDA Effect Size analysis
TSS: Total Sum Scaling normalization
FDR: False Discovery Rate
NCGS: non-celiac gluten sensitivity patients
GI: gastro intestinal
SCFA: short-chain fatty acids
FDR: First Degree Relative.
HC: Healthy Control.

## Declarations

### Ethics approval and consent to participate

“Not applicable”.

### Consent for publication

“Not applicable”.

### Authors’ contributions

Conceptualisation, L.J.M.-Z. and E.C.d.S.P.; methodology, J.A, and L.J.M.-Z.; writing—original draft preparation, J.A. and L.J.M.-Z; writing—critical review and editing, V. L-K, A.R.d.M.; funding acquisition, L.J.M.-Z. and E.C.d.S.P. All authors have read and agreed to the published version of the manuscript.

### Corresponding author

Correspondence to Enrique Carrillo de Santa Pau and Laura Judith Marcos-Zambrano

## Acknowledgements

Figure 1 and Figure 2 were designed with Biorender.com.

