## Supplementary material for "Metataxonomic review to elucidate the role of the microbiome in Celiac disease across the gastrointestinal tract": Suppl. material

**Figure S1.** Flowchart of the Study selection for literature review.

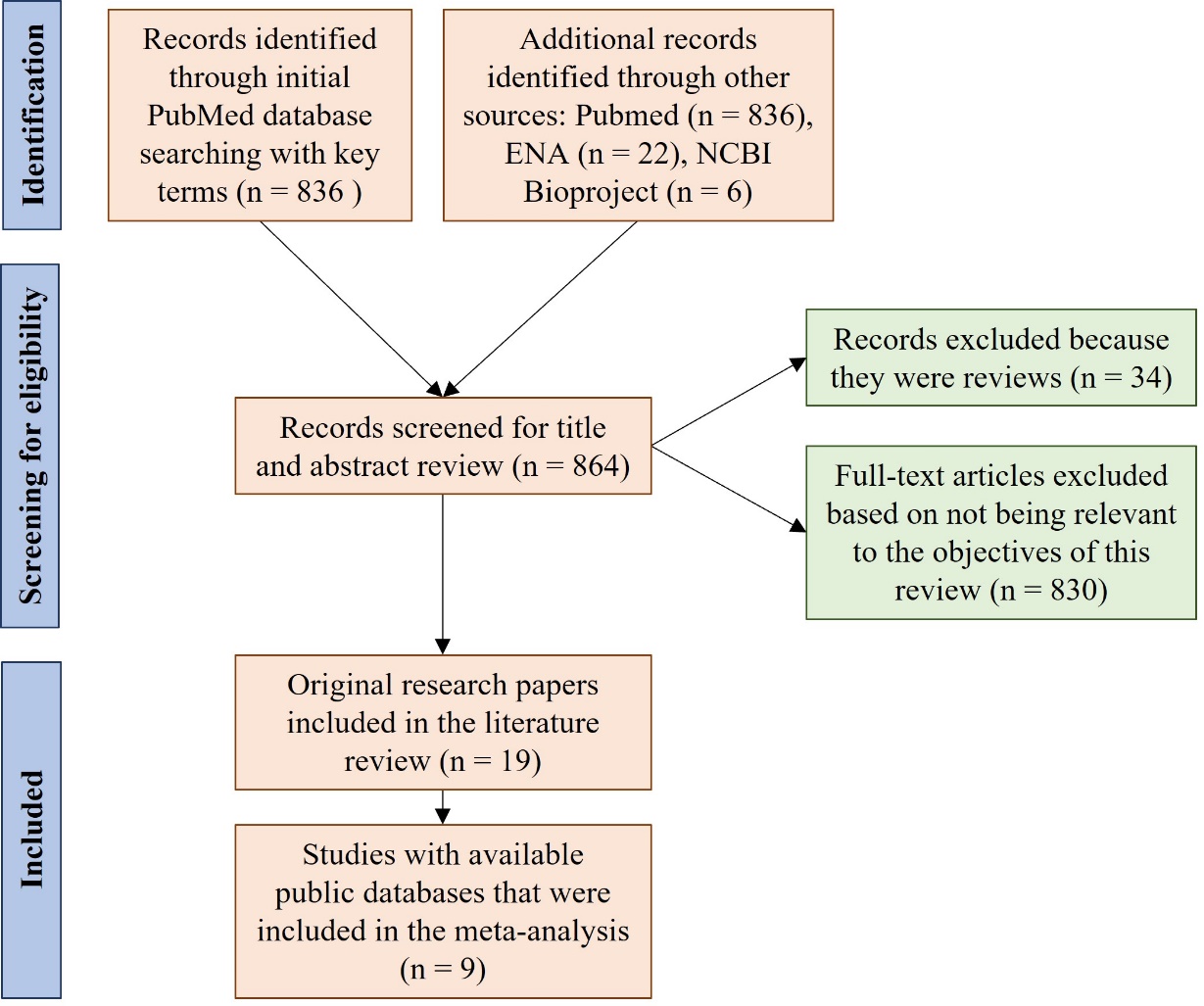

**Table S1.** Summary of the criteria used for the literature search and the statistics of the results.

| **Database** | **Target^1^** | **Terms** | **Total** | **New Selected^2^** | **Public data^3^** | **Other filters** |
| --- | --- | --- | --- | --- | --- | --- |
| **Pubmed** | Original research studies with newly generated data | (Celiac OR coeliac) AND (microb*) AND sequencing AND 16S | 21 | 13 | 5 | Full Text, datasets, journal articles, Humans, 2010-2020 |
| **Pubmed** | Original research studies with newly generated data | (Celiac OR coeliac) AND (microb*) AND 16S | 39 | 2 | 0 | Full Text, datasets, journal articles, Humans, 2010-2020 |
| **Pubmed** | Original research studies with newly generated data | (Celiac OR coeliac) AND (microb*) AND metagenomics | 49 | 0 | 0 | Full Text, datasets, journal articles, Humans, 2010-2020 |
| **Pubmed** | Original research studies with newly generated data | (Celiac OR coeliac) AND (microb*) | 727 | 0 | 0 | 2010-2020 |
| **ENA** | Original research studies with newly generated data | microb* AND (Celiac OR Coeliac) | 22 | 4 | 8 | NA |
| **NCBI Bioproject** | Original research studies with newly generated data | microb* AND (Celiac OR Coeliac) AND 16S | 6 | 0 | 5 | NA |
| **Pubmed** | Reviews | (microb*[Title]) AND (celiac[Title] OR coelic[Title]) | 17 | 17 | 0 | Full text, Review, Systematic Review, 2010-2020 |
| **Pubmed** | Reviews | (microb*) AND (Celiac OR coeliac) AND 16S AND Review | 1 | 1 | 0 | Full text, Review, Systematic Review, 2010-2020 |
| **Pubmed** | Reviews | (Celiac OR coeliac) AND (microb*) | 727 | 16 | 0 | 2010-2020 |

^1^ The type of paper intended to find. ^2^The additional studies found with certain terms, which were not found in the previous searches. ^3^The number of studies with data sets available in public databases.

**Table S2.** Main findings of selected papers after scoping review.

| **Group** | **Reference** | **Sample** | **Question Objective** | **Results** | **Methodology** | **Higthouputh method** | **Final conclusion** |
| --- | --- | --- | --- | --- | --- | --- | --- |
| **Children at risk of CeD** | Sellitto et al., 2012 | stool | How is the microbiome of infants at-risk before CeD onset?  May gut microbiota trajectory in early life be a predictor of celiac disease development?  Does time of first-time gluten exposure affect the time of CeD onset? | Microbiota of infants genetically at-risk for CeD was characterised by a low *Bacteroidetes* abundance and was highly different among children before 18 months of age but converged at 24 months, and the communities become similar to those of adults.  At 12 months of age more children in the group of early exposure to gluten presented antigliadin antibodies.  The children who developed CeD at 24 months of age belonged to the group of early exposure to gluten and showed a reduction in bacterial richness from 6 to 10 months. | Longitudinal study from birth to 24 months of age. Samples taken at 7 and 30 days, 6, 8, 10, 12, 18 and 24 months. No follow up.  **Patients:** 16 infants with first-degree relatives diagnosed with CeD that were positive for HLA DQ2 and/or HLA DQ8 genotypes.  **Diet:** Until 6 months of age breastfed. From 6 to 12 months: 8 children on GFD and the other 8 children on gluten containing diet. From 12 to 24 months all children on gluten containing diet | Pyrosequencing of the V1–V2 region of the 16S rRNA gene | No definitive answer about the predictivity of CeD by early microbiome composition can be drawn because only one children developed CeD and is not enough to make statistical analysis and achieve valid conclusions.  As the percentage of children with antigliadin antibodies in the group of early exposure to gluten was higher increasing the time of follow up to include CeD development at a later age would have been interesting. |
|  | Rintala et al., 2018 | stool | How is the microbiome of infants at-risk before CeD onset?  May gut microbiota trajectory in early life be a predictor of celiac disease development? | Microbiota’s alpha diversity in all at-risk children increased between 9 and 12 months of age.  Nine children (all girls) developed CeD at the median age of 3.5 years (range 2.6–4.2 years).  No differences among healthy controls and the group who develop CeD were detected.  No bias depending on the type of feeding (breast milk versus formula together or not with complementary feeding) was detected. | Longitudinal study from 9 to 12 months of age. Samples taken at 9 and 12 months of age. Five years follow up.  **Patients:** 27 infants with first-degree relatives diagnosed with CeD. All positive for the HLA DQ2.5 genotype.  **Diet:** The introduction of gluten on the diet was initiated from 4.4 to 5.7 months of age in all cases.  Similar to the early exposure group of Sellito et. al | Sequencing of the V4 region of the 16S rRNA gene using Illumina MiSeq platform | The hypothesis of at-risk children developiong CeD due to peculiarly vulnerable gut microbiota at early life is not supported by this study.  The individuals who develop CeD, do not already in the early infancy, have a distinct microbiota composition compared to other infants with risk-HLA-haplotype. |
|  | Olivares et al., 2018 | stool | How is the microbiome of infants at-risk before CeD onset?  May gut microbiota trajectory in early life be a predictor of celiac disease development? | 9 children developed CeD between 24 and 40 months and 1 at 82 months.  The diversity of microbiome increases among 4 and 6 months of age in children who remained healthy but not in those who develop CeD. Unlike Rintala et al. they did found changes among cases and controls.  No bias depending on the HLA-DQ, the type of feeding (breast milk versus formula together or not with complementary feeding), and the effect of antibiotic intake in the samples was found. | Longitudinal study from 4 to 6 months of age. Samples taken at 4 and 6 months of age  **Patients:** 20 infants with first-degree relatives diagnosed with CeD. Nineteen were positive for HLA DQ2 and/or HLA DQ8 genotypes. Five years follow up.  **Diet:** The introduction of gluten was initiated from 6 months of age in all cases. Similar to the early exposure group of Sellito et. al | Sequencing of the V1–V2 region of the 16S rRNA gene using Illumina MiSeq platform | Alterations in the early trajectory of gut microbiota in infants at CeD risk could influence predispose to CeD, although larger population studies are needed to confirm this hypothesis |
|  | Leonard et al., 2021 | stool | Do factors like HLA DQ2/DQ8 genotype, delivery mode, antibiotic exposure and infant feeding type impact on microbial composition of gut in at-risk infants?.  Unlike previous studies they did not evaluate predictive value of microbiota composition at an erly age in the onset of CeD. They did not knew the onset of the patients. | Genetic risk to develop CeD was associated with: Decreased abundance of *Streptococcus* and *Coprococcus* and with decreased abundance of *Veillonella*, *Parabacteroides* and *Clostridium perfringens* at 4-6 months of age. Increased abundance of *Bacteroides* and *Enterococcus* species at 0 months.  They also found differences in the microbiota regarding exposure to breastmilk and formula, and between antibiotic exposure and delivery mode. | Longitudinal study from birth to 6 months of age. Samples taken at 0 and 3 and 4-6months of age.  **Patients:**  31 infants with first-degree relatives diagnosed with CeD. 26 were positive for HLA DQ2 or HLA DQ8 genotype.  **Diet:** Study performed before the introduction of gluten in the diet was initiated. | Metagenome WGS Illumina MiSeq platform | Despite the study provides insights into taxonomic and functional shifts in the developing gut microbiota of infants at risk of CeD linking genetic and environmental risk factors to detrimental immunomodulatory and inflammatory effects., it is unclear whether they indeed contribute to the future development of CeD |
| **CeD in comparison with other groups of interest** | Cheng. et al., 2013 | duodenum | The main aim of the study was to characterise the total duodenal mucosal microbiota and evaluate the differences in pediatric CeD patients and healthy controls by using a high-throughput bacterial phylogenetic microarray (HITChip). | Authors do not find significant differences in the abundance of taxa between groups, neither at phylum nor at the genus level. Using Random Forest algorithm with a subpopulation of eight genus-like taxa, they were able to separate samples between CeD and healthy controls. | **Patients and diet:** Authors analysed the microbiome of 10 children between 3 and 14 years old, with newly diagnosed CeD, before GFD treatment, and nine children between 4 and 16 years old without CeD following an unrestricted gluten diet. | High-throughput bacterial phylogenetic microarray (HITChip).with oligos targeting the V1 and V6 regions of the16S rRNA gene | Bacteria from *Proteobacteria* were elevated in CeD and butyrate producers *Firmicutes* were diminished. Moreover, in CeD subjects, the increased expression of IL-10 and IFN-g may have partly resulted from the increased TLR9 expression and signalling. |
|  | Bodkhe et al., 2019 | duodenum and  stool | First-degree relatives (FDRs) of CeD patients may provide an opportunity to study gut microbiome in pre-disease state as FDRs are genetically susceptible to CeD? | Diversity measures were not significantly different between the disease condition (CeD), pre-disease (FDR) and control subjects. However, differences were observed at the level of amplicon sequence variant (ASV), suggesting alterations in specific ASVs between pre disease and diseased condition. Duodenal biopsies showed higher differences in ASVs compared to fecal samples indicating larger disruption of the microbiota at the disease site. | **Patients**: 23 patients with CeD (all of them were DQ2 and/or DQ8 positive), 15 healthy first-degree relatives (13 of them were DQ2 and/or DQ8 positive); and 24 non-related controls (6 of them were DQ2 and/or DQ8 positive).  **Diet**: Gluten-containing diet | Illumina MiSeq to sequence the V4 region of the 16S rRNA gene | The findings of the present study demonstrate differences in ASVs and predicts reduced ability of CeD faecal microbiota to degrade gluten compared to the FDR faecal microbiota. Further research is required to investigate the strain level and active functional profiles of FDR and CeD microbiota to better understand the role of gut microbiome in pathophysiology of CeD. |
|  | Pellegrini et al., 2017 | duodenum | The objective of this study was to evaluate the gut inflammatory profile and microbiota in patients with T1D compared with healthy control (CTRL) subjects and patients with celiac disease (CD) as gut inflammatory disease controls. | The T1D duodenal mucosal microbiome results were different from the other groups, with an increase in *Firmicutes* and *Firmicutes/Bacteroidetes* ratio and a reduction in *Proteobacteria* and *Bacteroidetes*. The expression of genes specific for T1D inflammation was associated with the abundance of specific bacteria in the duodenum. | The inflammatory status and microbiome composition were evaluated in biopsies of the duodenal mucosa.  **Patients:** Patients with T1D (n = 19), in patients with CD (n = 19), and CTRL subjects (n = 16).  Diet: | Pyrosequencing of the V3- V5 region of the 16S rRNA gene | Authors showed that CeD patients share some similitudes in the microbial profile with T1D i.e., reduction in *Bacteroidetes* phylum and the class *Clostridia*, but the microbial profile differed according to the abundance of *Proteobacteria*, being increased in CeD, particularly *Gammaproteobacteria*. Moreover, the study shows that duodenal mucosa in T1D presents disease-specific abnormalities in the inflammatory profile and microbiota. |
|  | Garcia-Mazcorro et al., 2018 | duodenum and stool | How is the gut microbiota in Mexican people afflicted with Gluten-related disorders (GRDs)? | The genus *Actinobacillus* and the family *Ruminococcaceae* were higher in the duodenal and faecal microbiota of NCGS patients, respectively, while *Novispirillum* was higher in the duodenum of CD patients. | **Patients and diet:** Authors characterise the duodenal and faecal microbial communities of six patients with CeD on a gluten-containing diet (between 25 and 73 years old), twelve non-CeD controls (between 23 and 64 years old) and twelve Non-Celiac Gluten Sensitivity (NCGS) patients (between 21 and 59 years old). | Illumina MiSeq to sequence the V4 region of the 16S rRNA gene | This study generates valuable preliminary data about the relationship between the gut microbiota and gluten-related disorders in Mexican people. Interestingly, the four-week consumption of GFD was associated with an increased abundance of *Pseudomonas* in duodenal biopsies of patients with these disorders, particularly in NCGS patients. This change was noticed despite a general lack of differences in richness or diversity. |
|  | Panelli et al., 2020 | duodenum, stool, and saliva | Authors performed a comparative analysis of the gut microbiota in adulthood CD to evaluate whether: (i) dysbiosis anticipates mucosal lesions, (ii) gluten-free diet restores eubiosis, (iii) refractory CD has a peculiar microbial signature, and (iv) salivary and fecal communities overlap the mucosal one | A reduction of both alfa and beta diversity in CD, already evident in the potential form and achieving nadir in refractory CD, was evident. Taxonomically, mucosa displayed a significant abundance of *Proteobacteria* and an expansion of *Neisseria*, especially in active patients, while treated celiacs showed an intermediate profile between active disease and controls. The saliva community mirrored the mucosal one better than stool. | Microbial communities in the duodenal, salivary and faecal samples of 83 individual's classified in four groups depending on their disease state, their type of diet, and the presence of villous atrophy in their duodenal biopsy.  **Patients and diet:** Individuals with CeD were classified into three groups. 1. Active (ACD), those on a gluten-containing diet presenting villous atrophy; 2. Refractory (RCD), those on a GFD presenting villous atrophy and; 3. Treated (TCD), those on a gluten-free without the presence of villous atrophy. Non-CeD individuals were classified into two groups. 1. Potential (PCD) and controls (C), healthy individuals on a gluten-containing diet and; controls, individuals with functional dyspepsia on a gluten-containing diet. All ACD and RCD were HLADQ2 and/or HLADQ8 positive. Some TCD, PCD and C were HLADQ2 and/or HLADQ8 positive. | Illumina MiSeq to sequence the V3-V4 region of the 16S rRNA gene | Expansion of pathobiontic species anticipates villous atrophy and achieves the maximal divergence from controls in refractory CD. Gluten-free diet results in incomplete recovery. The overlapping results between mucosal and salivary samples indicate the possible use of saliva as a diagnostic fluid. |
| GFD effect on the microbiome of CeD, NCGS patients and of healthy people and of healthy controls | Bonder et al., 2016 | stool | The main aim of the study was to investigate the diet-related changes on the level of taxonomic units as on the predicted bacterial pathways. | Inter-individual variation in the gut microbiota remained stable during this short-term GFD intervention. A number of taxon-specific differences were seen during the GFD: the most striking shift was seen for the family *Veillonellaceae* (class *Clostridia*), which was significantly reduced during the intervention Seven other taxa also showed significant changes; the majority of them are known to play a role in starch metabolism. Authors also find stronger differences in pathway activities: 21 predicted pathway activity scores showed significant association to the change in diet. | GFD study for 13 weeks on 21 aged between 16 and 61 years (mean age, 36.3 years) without any known food intolerance. Measurements started at T=0 when patients were in Gluten-containing diet; they then follow a GFD for 4 weeks and a "wash-out" period of five weeks. Finally, they return to their habitual diets (HD, gluten-containing) for four weeks (T = 5–8). They collected stool at all points and blood ad T6 and T8 | 454 pyrosequencing of the V3-V4 region of the 16S rRNA gene | A GFD changes the gut microbiome composition and alters the activity of microbial pathways. |
|  | Ercolini et al.,2016 | saliva | How is the salivary microbiota and metabolome of Saharawi celiac children in response to the change from the traditional African- to the Italian-style gluten-free diet?. | An Italian-style gluten-free diet caused increases in the abundance of *Granulicatella*, *Porphyromonas* and *Neisseria* and decreases in *Clostridium*, *Prevotella* and *Veillonella*, altering the ‘salivary type’ of the individuals. Furthermore, operational taxonomic unit co-occurrence/exclusion patterns indicated that the initial equilibrium of co-occurring microbial species was perturbed by a change in diet: the microbial diversity was reduced, with a few species out-competing the previously established microbiota and becoming dominant. Analysis of predicted metagenomes revealed a remarkable change in the metabolic potential of the microbiota following the diet change, with increased potential for amino acid, vitamin and co-factor metabolism. High concentrations of acetone and 2-butanone during treatment with the Italian-style gluten-free diet suggested metabolic dysfunction in the Saharawi celiac children. | **Patients and diet:** Authors studied the influence of a change on GFD over the microbiome of fourteen celiac children (8.4 +/- 0.7 years old) following an African- style GFD during two years before the 60 days of treatment with an Italian-style gluten-free diet. African diet consisted of gluten-free cereals, legumes, vegetables, high carbohydrate content, fibre and non-minimal protein contents. In contrast, the Italian diet was based on animal proteins, sugars, starch, fats and fibre. | 16S rRNA gene region V1-V3 was sequenced by using 454 pyrosequencing. | The findings of this study support the need for a translational medicine pipeline to examine interactions between food and microbiota when evaluating human development, nutritional needs and the impact and consequences of westernisation. |
|  | D'Argenio. et al., 2016 | duodenum | The aim of this study was to determine the CeD-associated duodenal dysbiosis in adult celiac patients and elucidate the mechanisms influencing in CeD development or exacerbation. . | *Proteobacteria* was the most abundant and *Firmicutes* and *Actinobacteria* the least abundant phyla in the microbiome profiles of active CeD patients. Members of the *Neisseria* genus (*Betaproteobacteria* class) were significantly more abundant in active CeD patients than in the other two groups ( *P* =0.03). *Neisseria* *flavescens* (CD- Nf ) was the most abundant *Neisseria* species in active CeD duodenum. Whole-genome sequencing of CeD- Nf and control- Nf showed genetic diversity of the iron acquisition systems and of some hemoglobin-related genes. CeD- Nf was able to escape the lysosomal compartment in CaCo-2 cells and to induce an infl ammatory response. | **Patients and diet:** Authors characterise the duodenal microbiome communities of 41 individuals, 20 with CeD on a gluten-containing diet (mean age 38±12 years old), six with CeD on a GFD since two years before the study (mean age 39±11years old) and the control group constituted by seven patients (mean age 42±16 years old). | Pyrosequencing of the V4–V6 region of the 16S rRNA gene | Marked dysbiosis and an abundance of a peculiar CeD- Nf strain characterize the duodenal microbiome in active CeD patients thus suggesting that the CeD-associated microbiota could contribute to the many inflammatory signals in this disorder. |
|  | Nistal. et al., 2016 | duodenum | How is the composition of the duodenal microbiota between CeD patients and non‐CeD controls? | The sequences analysis showed that the majority of the reads were classified within two phyla: *Firmicutes* and *Proteobacteria*. Bacterial richness and diversity were higher in non‐CeD controls than in untreated CeD patients, but the differences were not statistically significant. The principal coordinates analysis revealed that bacterial communities of non‐CeD controls and untreated CeD patients were dispersed without forming a clear group according to diagnosis of CeD. | **Patients and diet:** Authors characterise the duodenal microbiome communities of eighteen adults, nine CeD on a gluten-containing diet and nine non-CeD controls. | Pyrosequencing of the V4 region of the 16S rRNA gene | There were no statistically significant differences in the upper small intestinal composition of bacterial communities between untreated CeD patients and non‐CeD controls. |
| **Upper gastrointestinal tract microbiome in CeD.** | Francavilla et al., 2014 | saliva | This study aimed to investigate the salivary microbiota and metabolome of 13 children with celiac disease (CeD) under a gluten-free diet (treated celiac disease [T-CeD]). The same number of healthy children (HC) was used as controls. | Compared to HC, the number of some cultivable bacterial groups (e.g., total anaerobes) significantly (P < 0.05) differed in the saliva samples of the T-CeD children. Pyrosequencing data showed the highest richness estimator and diversity index values for HC. Levels of *Lachnospiraceae*, *Gemellaceae*, and *Streptococcus sanguinis* were highest for the T-CeD children. *Streptococcus thermophilus* levels were markedly decreased in T-CeD children. The saliva of T-CeD children showed the largest amount of *Bacteroidetes* together with the smallest amount of *Actinobacteria*. As shown by multivariate statistical analyses, the levels of organic volatile compounds markedly differentiated T-CeD children. Some compounds (e.g., ethyl-acetate, nonanal, and 2-hexanone) were found to be associated with T-CeD children. | The salivary microbiota was analysed by an integrated approach using culture-dependent and -independent methods. Metabolome analysis was carried out by gas chromatography-mass spectrometry–solid-phase microextraction.  **Patients and diet:** compared the salivary microbiome of children (median age, 10 +/- 1.4 years) with CeD treated with GFD for at least two years and healthy control children without CeD or any other known food intolerances who followed an unrestricted diet. | Pyrosequencing of the amplified V1–V3 region of the 16S rRNA gene by 454 technology | CeD is associated with oral dysbiosis that could affect the oral metabolome. |
|  | Tian et al., 2017 | saliva | Are that enzymes produced by oral bacteria involved in gluten processing in the intestine and susceptibility to celiac disease? | Salivary glutenase activities were higher in CD patients compared to controls, both before and after normalization for protein concentration or bacterial load. The oral microbiomes of CD and RCD patients showed significant differences from that of healthy subjects, e.g., higher salivary levels of lactobacilli (P 0.05), which may partly explain the observed higher gluten-degrading activities. | They sequenced the salivary microbiome of 21 CeD patients responding to a GFD (for an average of 43 months), eight refractory CeD (R-CeD) patients on a GFD for an average of 85 months who had persistent GI symptoms. Twenty healthy subjects without CeD or gluten sensitivity and twelve patients without CeD reporting functional GI complaints.  Stimulated whole saliva was collected from patients with CD in remission (n 21) and refractory CD (RCD; n 8) and was compared to healthy controls (HC; n 20) and subjects with functional GI complaints (n 12). Salivary gluten-degrading activities were monitored with the tripeptide substrate Z-Tyr-Pro-Gln-pNA and the -gliadin-derived immunogenic 33- mer peptide. The oral microbiome was profiled by 16S rRNA-based MiSeq analysis. | Sequencing of the V3 to V4 hypervariable region of the 16S rRNA gene using MiSeq (Illumina) technology. | While the pathophysiological link between the oral and gut microbiomes in CD needs further exploration, the presented data suggest that oral microbe-derived enzyme activities are elevated in subjects with CD, which may impact gluten processing and the presentation of immunogenic gluten epitopes to the immune system in the small intestine. |
|  | Iaffaldano et al., 2018 | Oropharynx exudate | Authors investigate the oropharyngeal microbiome in CD patientsand controls to evaluate whether this niche share microbial composition with the duodenum | *Bacteroidetes*, *Proteobacteria* and *Firmicutes* differed significantly between the three groups. In particular, *Proteobacteria* abounded in a-CD and *Neisseria* species mostly accounted for this abundance (p < 0.001), whereas Bacteroidetes were more present in control and GFD microbiomes. Microbial functions prediction indicated a greater metabolic potential for degradation of aminoacids, lipids and ketone bodies in a-CD microbiome than in control and GFD microbiomes, in which polysaccharide metabolism predominated. | **Patients and diet**: Symptomatic CeD on an unrestricted diet (a-CeD), CeD patients on GFD (CeD-GFD) and healthy control subjects.  We characterized by 16S rRNA gene sequencing the oropharyngeal microbiome in 14 a-CD, 22 GFD patients and 20 controls. | V4-V6 region of the 16S rRNA | Results suggest a continuum of a-CD microbial composition from mouth to duodenum. Authors speculate that microbiome characterization in the oropharynx, which is a less invasive sampling than the duodenum, could contribute to investigate the role of dysbiosis in CD pathogenesis. |
| **Dietary interventions, prebiotics and hookworm infections** | Quagliariello et al., 2016 | stool | This work is aimed at the assessment of the impact of the administration of two *Bifidobacterium breve* strains on the gut microbiota composition of coeliac patients compliant to a GFD and, at the same time,it evaluates the difference in the intestinal colonization of coeliac subjects on a GFD for several years with respect to healthy subjects. | The comparison between CD subjects and Control group revealed an alteration in the intestinal microbial composition of coeliacs mainly characterized by a reduction of the Firmicutes/Bacteroidetes ratio, of Actinobacteria and Euryarchaeota. Regarding the effects of the probiotic, an increase of Actinobacteria was found as well as a re-establishment of the physiological Firmicutes/Bacteroidetes ratio. | They selected 40 CeD (positive for serological markers and biopsy analysis) between 1 and 19 years old and 16 healthy children. Microbial DNA was extracted from faeces of 40 coeliac children before and after probiotic or placebo administration and 16 healthy children (Control group). Sequencing of the amplified V3-V4 hypervariable region of 16S rRNA gene as well as qPCR of Bidobacterium spp., Lactobacillus spp., Bacteroides fragilis group Clostridium sensu stricto and enterobacteria were performed. | V3-V4 hypervariable region of 16S rRNA gene as well as qPCR | Therefore, a three-month administration of *B. breve* strains helps in restoring the healthy percentage of main microbial components. |
|  | Giacomin et al., 2016 | duodenum | In the present investigation, we have assessed, for the first time, the changes in the microbiota at the site of infection by a parasitic helminth (hookworm) and gluten-dependent inflammation in humans with CeD using biopsy tissue from the duodenum | Hookworm infection and gluten exposure were associated with an increased abundance of species within the Bacteroides phylum, as well as increases in the richness and diversity of the tissue-resident microbiota within the intestine, results that are consistent with previous reports using other helminth species in humans and animal models | The authors included six treated CeD patients (HLA-DQ2+ or HLA-DQ8+) on a strict GFD (> 5 years) who were infected with *N. americanus* (Trial subjects) and then underwent exposure to escalating doses of dietary gluten, with a 10–50 mg/day micro-challenge from weeks 12–24, followed by intermittent twice-weekly 1 g/day gluten challenge from weeks 24 to 36 (approximately 350 mg gluten/day). Before experimental infection (T0), as well as at 24 (T24) and 36 weeks (T36) post-infection microbiome of duodenal samples was sequenced. Besides, individual duodenal samples from six hookworm-naïve volunteers with active CeD (diagnosed as Marsh grade 3) (Control subjects) were also included for comparative purposes. | High-throughput sequencing of the V3-V4 hypervariable region of the bacterial 16S rRNA gene was performed on an Illumina MiSeq platform | this may represent a mechanism by which parasitic helminths may restore intestinal immune homeostasis and exert a therapeutic benefit in CeD, and potentially other inflammatory disorders. |

**Table S3.** Results of the analysis of differential abundance of metabolic pathways. Only the differentially abundant pathways are shown in a statistically significant way. Positive values represent an increase in gene abundance in controls compared to cases and negative values represent an increase in gene abundance in cases compared to controls.

| **Pathway** | **Tissue** | **log_2_FC*** | ***P* Value** | **FDR**** | **Controls** |
| --- | --- | --- | --- | --- | --- |
| Degradation of D-glucarate I | duodenum | -4,03 | 3,20E-15 | 1,03E-12 | Disminished |
| Degradation of L-arabinose IV | duodenum | -3,93 | 5,12E-10 | 2,75E-08 | Disminished |
| Degradation of D-glucarate and D- galactarate | duodenum | -3,89 | 1,47E-14 | 1,58E-12 | Disminished |
| Degradation D-galactarate I | duodenum | -3,84 | 1,47E-14 | 1,58E-12 | Disminished |
| Degradation of biogenic amines | duodenum | -3,22 | 2,94E-11 | 2,38E-09 | Disminished |
| Degradation of lactose and galactose I | duodenum | 3,04 | 3,02E-09 | 1,08E-07 | Disminished |
| Fermentation of hexitol to lactate, formate, ethanol and acetate | duodenum | 3,29 | 4,36E-11 | 2,81E-09 | Disminished |
| Biosynthesis of ADP-L-glycero-β-D-manno-heptose | stool | -5,4039 | 6,60E-27 | 1,83E-24 | Disminished |
| Fermentation of acetyl-CoA to butanoate II | stool | -4,6659 | 1,36E-18 | 2,15E-17 | Disminished |
| Thiamine diphosphate II biosynthesis | stool | -4,4658 | 2,89E-22 | 7,69E-21 | Disminished |
| Biosynthesis of the thiazole component of thiamine diphosphate | stool | -4,3899 | 1,39E-19 | 2,75E-18 | Disminished |
| (KDO) 2-lipid IVA transferase III (Chlamydia) | stool | -3,5326 | 5,35E-23 | 2,12E-21 | Disminished |
| CMP-3-deoxy-D-manno-octulosonate biosynthesis | stool | -3,4348 | 1,11E-22 | 3,86E-21 | Disminished |
| Biosynthesis of lipid IVA | stool | -3,4288 | 3,04E-22 | 7,69E-21 | Disminished |
| Guanocine III nucleotide degradation | stool | -3,4273 | 3,28E-18 | 4,79E-17 | Disminished |
| Thiamine diphosphate I biosynthesis | stool | -3,3263 | 1,68E-24 | 1,56E-22 | Disminished |
| Biosynthesis of preQ0 | stool | -3,2554 | 7,14E-25 | 9,92E-23 | Disminished |
| Gluconeogenesis I | stool | -3,1878 | 1,91E-23 | 1,06E-21 | Disminished |
| Nitrate VI reduction (assimilation) | stool | -3,1798 | 1,91E-23 | 1,06E-21 | Disminished |
| Tetrahydrofolate biosynthesis | stool | -3,0552 | 3,68E-14 | 3,79E-13 | Disminished |
| Queuosin biosynthesis (de novo) | stool | -3,0297 | 1,25E-23 | 8,68E-22 | Disminished |
| Synthesis of peptidoglycan II | saliva | -5,4509 | 3,53E-08 | 9,86E-06 | Disminished |
| Degradation of protocatechuate II | pharynx | -9,2733 | 2,40E-09 | 2,85E-07 | Increased |
| Degradation of aromatic compounds by β-ketoadipate | pharynx | 8,6482 | 4,08E-09 | 2,85E-07 | Increased |
| Catechol III degradation | pharynx | 8,6482 | 4,12E-09 | 2,85E-07 | Increased |
| superpathway of 2,3-butanediol I biosynthesis | pharynx | 8,5633 | 4,56E-09 | 2,85E-07 | Increased |
| Catechol degradation to β-ketoadipate | pharynx | 8,4808 | 4,92E-09 | 2,85E-07 | Increased |
| Degradation of L-leucine I | pharynx | 8,3194 | 1,05E-08 | 5,04E-07 | Increased |
| superpathway of 2,3-butanediol II biosynthesis | pharynx | 9,2733 | 2,40E-09 | 2,85E-07 | Increased |
| Catechol degradation to β-ketoadipate | pharynx | 8,6482 | 4,08E-09 | 2,85E-07 | Increased |

* log2 of the Fold Change (FC); that is, log2 of the ratio between the relative abundance of a taxon between cases and controls.

** False discovery rate or p-value adjusted for multiple comparisons using the Benjamini-Hochberg test
